# Morphogenesis of bacterial colonies in polymeric environments

**DOI:** 10.1101/2024.04.18.590088

**Authors:** Sebastian Gonzalez La Corte, Corey A. Stevens, Gerardo Cárcamo-Oyarce, Katharina Ribbeck, Ned S. Wingreen, Sujit S. Datta

**Author notes:** **Corresponding authors** Correspondence and requests for materials should be addressed to Sujit S. Datta and Ned S. Wingreen.

## Abstract

Many bacteria live in polymeric fluids, such as mucus, environmental polysaccharides, and extracellular polymers in biofilms. However, lab studies typically focus on cells in polymer-free fluids. Here, we show that interactions with polymers shape a fundamental feature of bacterial life—how they proliferate in space in multicellular colonies. Using experiments, we find that when polymer is sufficiently concentrated, cells generically and reversibly form large serpentine “cables” as they proliferate. By combining experiments with biophysical theory and simulations, we demonstrate that this distinctive form of colony morphogenesis arises from an interplay between polymer-induced entropic attraction between neighboring cells and their hindered ability to diffusely separate from each other in a viscous polymer solution. Our work thus reveals a pivotal role of polymers in sculpting proliferating bacterial colonies, with implications for how they interact with hosts and with the natural environment, and uncovers quantitative principles governing colony morphogenesis in such complex environments.

## 1 Introduction

Many bacteria live in polymeric fluids, such as mucus that lines the airways, gut, and cervico-vaginal tract in the body [1, 2], exopolymers in the ocean [3], and cell-secreted extracellular polymeric substances (EPS) that encapsulate biofilms [4]. However, lab studies of bacteria typically focus on cells in polymer-free fluids. As a result, despite their prevalence, how extracellular polymers influence bacterial behavior remains poorly understood.

Recent work hints that interactions with polymers can dramatically alter how individual motile cells swim [5–8] and aggregate with other cells [9–13]. Nevertheless, the possible influence of polymers on another fundamental characteristic of bacterial life—spatial proliferation in a multicellular colony [14, 15]—remains unknown. Colony morphology can be biologically important. For example, studies of bacteria proliferating on planar surfaces show that their resulting colonies can exhibit a variety of morphologies [14, 16–25], which in turn can affect cell-cell signaling [26], genetic diversity [27–30], and colony resilience and susceptibility to external stressors [9, 31, 32]. Hence, we ask: Do interactions with polymers sculpt the morphology of proliferating bacterial colonies?

Here, we demonstrate that this is indeed the case, and we elucidate the underlying mechanisms. In nature, many bacteria are non-motile or lose motility [33–56], but still continue to proliferate in colonies; indeed, the loss of flagella is often associated with pathogenesis and bacterial adaptation to diseased mucosal environments, such as in cystic fibrosis. Therefore, we focus on the spatial proliferation of rod-shaped, non-motile mutant cells that lack flagella. By performing experiments with these bacteria in polymer solutions, we find that when polymer is sufficiently concentrated, cells form large-scale “cables” as they proliferate in a colony—in stark contrast to forming a random dispersion, as in the conventionally studied polymer-free case. This characteristic cable morphology arises independent of variations in cell type and polymer composition across three different species of bacteria and seven different polymers, including mucins, a key component of mucus in the body. By combining experiments, theoretical modeling, and agent-based simulations, we trace the origin of cable formation to an interplay between polymer-induced entropic attraction between neighboring cells and their hindered ability to diffusively separate from each other after growth and division in a viscous polymer solution. Our work thus reveals a pivotal role of polymers in shaping proliferating bacterial colonies and provides quantitative principles to predict and control these morphodynamics more broadly.

## 2 Results

### Non-motile bacteria proliferating in polymer solutions form long multicellular cables, across different species and solution compositions

We use confocal microscopy to directly visualize non-motile bacteria, constitutively expressing green fluorescent protein (GFP) in their cytoplasms, as they proliferate in nutrient-rich solutions with polymers added at a defined mass concentration *c*. To start, we use *Escherichia coli* as a model bacterium and mucus obtained from human primary transverse colon cells [57] as the polymer solution. In polymer-free solution, the cells continually grow, divide, separate from each other, and thermally diffuse apart, eventually forming a random dispersion (Figure 1a, Movie S1). Colony morphogenesis is completely different in the mucus solution: cells pack side-by-side up to a limited bundle size and then remain oriented end-to-end for subsequent divisions, eventually forming an intertwined network of long, serpentine, multicellular cables (Fig. 1b, Movie S2). This phenomenon is not dependent on the source of the mucus: we observe similar colony morphologies using purified native porcine intestinal mucin (Muc2) at *c* = 0.5 w*/*v% (Fig. S6, Movie S3), which is also representative of human intestinal mucus [1, 2].

**Fig. 1:**
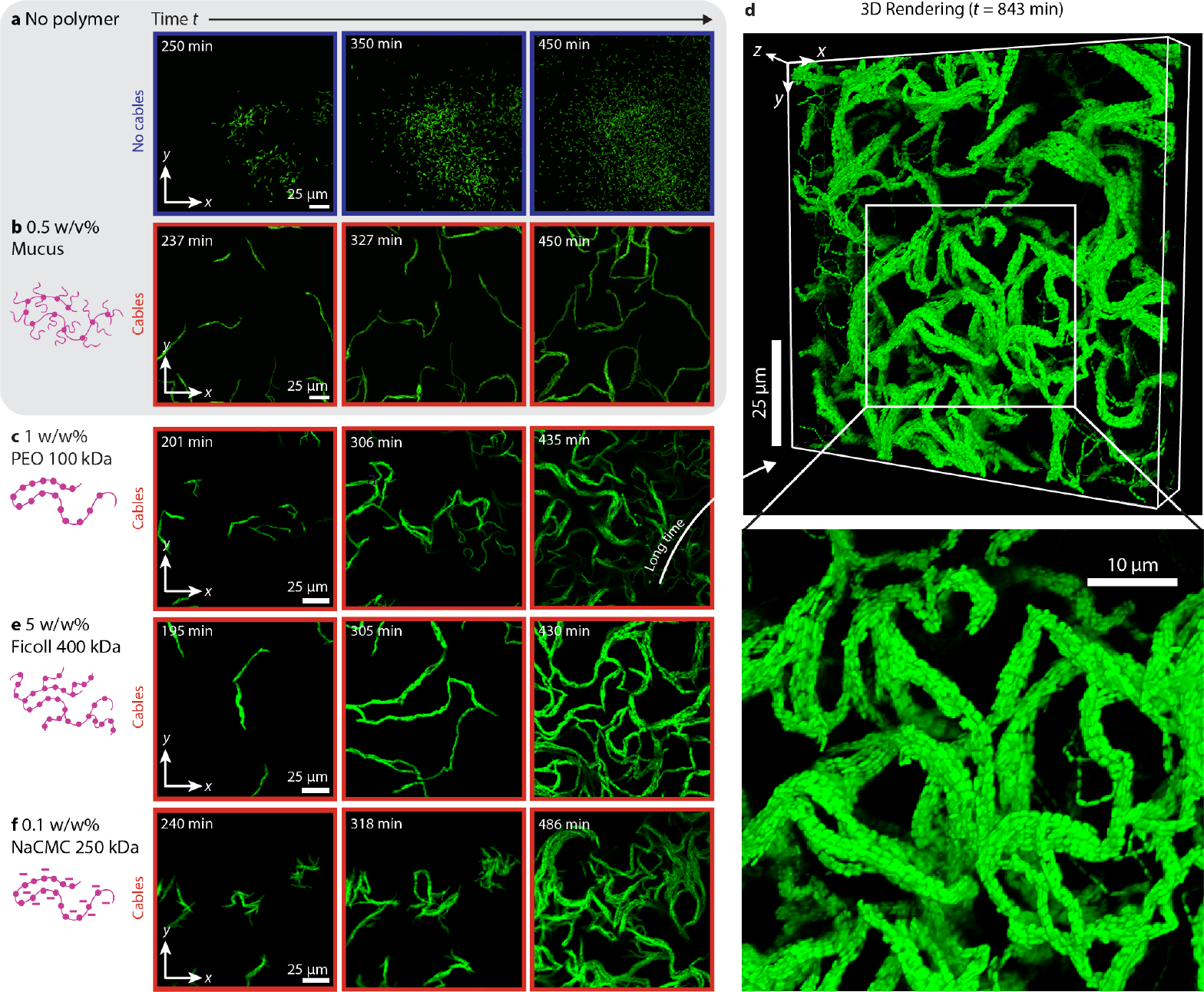
Non-motile bacteria generically form cables as they proliferate in polymer solutions. **a**, Time sequence of non-motile *E. coli* proliferating in nutrient-rich polymer-free fluid; the cells form a random dispersion. **b–c, e–f**, Same as in **a**, but with added **b** Mucus (0.5 w/v%), **c** PEO 100 kDa (1 w/w%), **e** Ficoll 400 kDa (5 w/w%), or **f** NaCMC 250 kDa (0.1 w/w%); in these polymeric fluids, cells form large, serpentine cables. Full dynamics can be seen in Movies S1–6. **d**, Three-dimensional rendering of a small volume of the colony shown in **c** after 843 min from the initiation of imaging, showing the internal structure of cables.

To explore the generality of cable formation, we repeat this experiment, but replacing the mucins with 100 kDa polyethylene oxide (PEO), a chemically-inert, uncharged, non-adsorbing, synthetic polymer [58–61]. For sufficiently concentrated PEO solutions, we again observe cable formation, as exemplified by the confocal micrographs in Fig. 1c and Movie S4, indicating that cable formation does not require specific biochemical interactions with mucins [58, 62, 63]. A 3D reconstruction of the final colony is shown in Fig. 1d, illustrating that the transverse cross-section of each cable has multiple cells packed side-by-side, all nematically aligned along their long axis. This network of cables extends into the bulk of the sample away from the solid boundaries, and similar cable morphologies arise for continuously-rotated samples for which cells do not settle, but remain in the bulk (Fig. S7)—indicating that cable formation is not strongly influenced by the presence of boundaries in the experiments.

To further test the generality of cable formation, we repeat this experiment, but using an even broader range of different polymers: PEO of three different molecular weights; Ficoll, a highly-branched, uncharged polysaccharide; and sodium carboxymethyl cellulose (NaCMC), a linear, anionic polysaccharide. In all cases, we observe cable formation when the polymer solution is sufficiently concentrated, as exemplified by Fig. 1e–f and Movies S5–6. Thus, cable formation arises independent of variations in polymer chemistry, molecular weight, molecular architecture, and charge. As a final test of the generality of this phenomenon, we perform the same experiment using non-motile strains of two other species of bacteria that also inhabit polymeric environments: *Vibrio cholerae*, a gut pathogen, and *Pseudomonas aeruginosa*, a lung pathogen. We again observe cable formation in both cases (Fig. S8), indicating that this phenomenon arises across different cell types.

Taken together, these results demonstrate that non-motile bacterial colonies proliferating in sufficiently-concentrated polymer solutions generically form large, serpentine, multicellular cables.

### Polymer-induced entropic attraction between cells is required for cable formation

Why do cables form? Given that cable formation does not require specific biochemical interactions, we hypothesize that it instead arises from some other physicochemical influence of polymers on cells. For example, could it be that when polymers are sufficiently concentrated, they form a meshlike network that entraps the individual cells, retaining them in the end-to-end configuration after division and thereby promoting cable formation? Our data indicate that this is not the case: across all the different conditions tested, we find no correlation between the threshold polymer concentration at which cables begin to form and *c*^∗^, the concentration at which the different polymer molecules begin to overlap and form an interconnected network (Table S1). Similarly, we find no correlation between the onset of cable formation and the solution viscosity *η* (Table S1), indicating that cable formation does not arise solely because of the reduced ability of cells to diffusively separate from each other post-division in a viscous polymer solution.

Another possibility is that the osmotic pressure Π_osm_ exerted by the polymers alters how the individual cells grow, somehow causing them to proliferate in cables. As shown in Table S1, however, cable formation arises at osmotic pressures as low as ∼10s of Pascals — far lower than the pressure at which bacterial physiology is typically altered, ∼100 kPa [64–70], arguing against this possibility. Indeed, growth curves measured for shaken cultures show no appreciable differences upon polymer addition (Fig. S9). Moreover, the osmotic pressure at which cables begin to form varies across different polymers (Table S1), indicating that osmotic pressure does not solely control cable formation.

However, the osmotic pressure of a polymer solution can influence proliferating cells in another way: by inducing attractive interactions between them. Consider two microscale particles—e.g., adjacent cells (green in Fig. 2a)—in a polymer solution (purple). The centers of the surrounding polymer chains are excluded from a small region surrounding each particle (light grey), referred to as its excluded volume. When the particles are close enough to each other, their excluded volumes overlap and surrounding polymers are depleted from this overlapping region of volume *V*_ev_ (dark grey). This depletion of polymers gives rise to an osmotic pressure difference Π_osm_ across the particles, pushing them together, as schematized in Fig. 2a(i). It is well documented in colloidal science that this entropic effect forces passive, non-proliferating particles to stick together, and can be modeled as an attractive “depletion” interaction between pairs of particles. The magnitude of this depletion interaction energy is *U*_dep_ = Π_osm_*V*_ev_ [71, 72], where Π_osm_ is determined by the polymer size and concentration [73] and *V*_ev_ is also determined by the polymer size and concentration, as well as the particle sizes, shapes, and orientations [74]. Could this polymer-induced depletion attraction between cells somehow cause them to proliferate in cables?

**Fig. 2:**
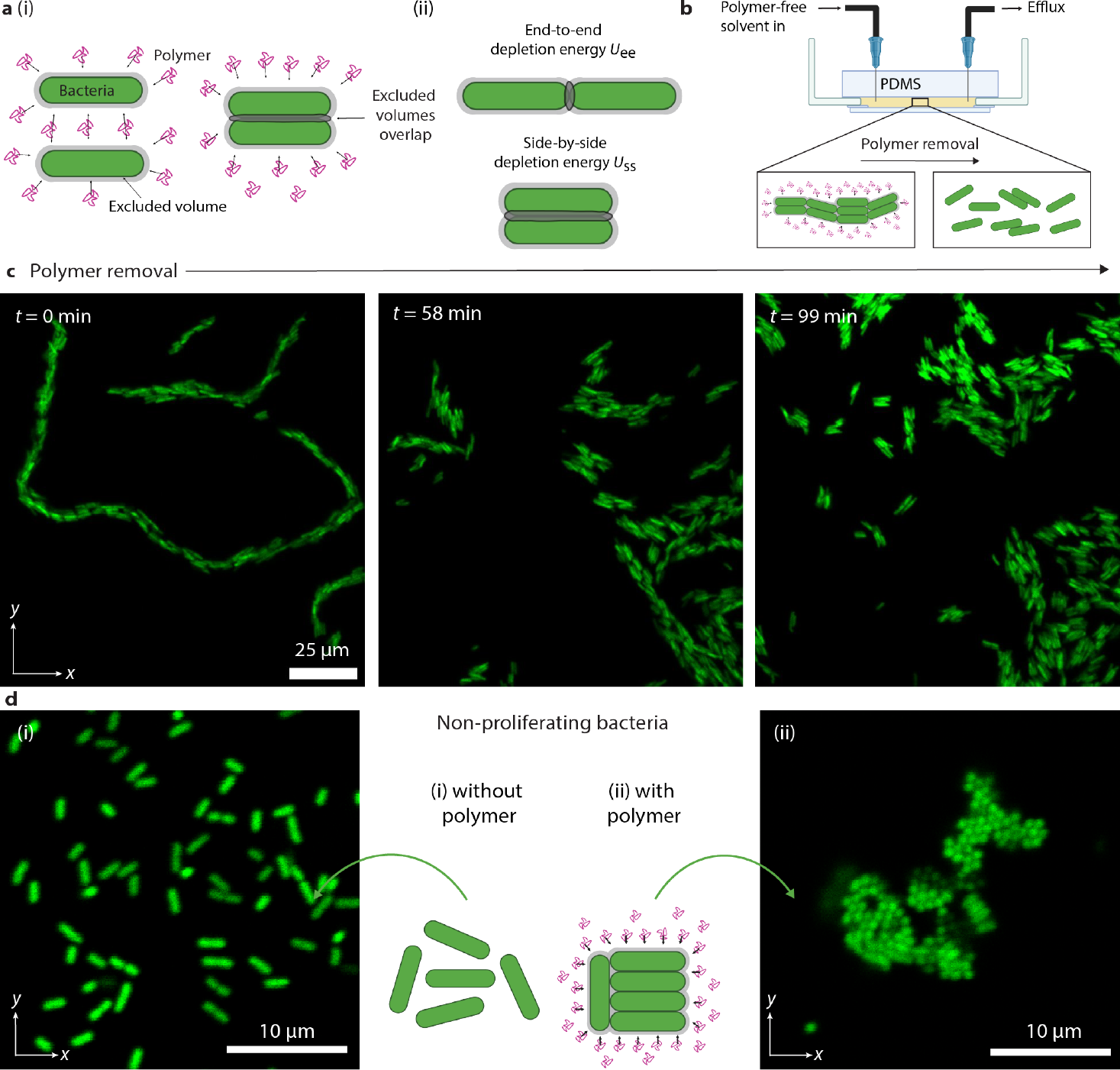
Polymer-induced entropic attraction between cells is required for cable formation. **a**, (i) Schematic of polymer (pink)-induced attraction between cells (green); the shaded grey regions show the excluded volume around each cell that is inaccessible to the centers of the polymer chains. As cells approach each other, their excluded volumes overlap (dark shaded regions), causing polymer to be depleted from the interstitial space. The osmotic pressure exerted by the surrounding polymers (arrows) drives the cells together, thereby resulting in attraction between cells. (ii) The overlapping excluded volume, and therefore the magnitude of the depletion interaction potential, is smaller for cells arranged end-to-end (top) versus side-by-side (bottom). **b**, Schematic of experiments in which cables formed in polymeric fluid are gently flushed with polymer-free fluid. **c**, Time sequence showing cables disintegrating when polymers are removed from their surroundings; polymer-free solvent is introduced at *t* = 0. Full dynamics can be seen in Movie S7. **d**, Non-proliferating cells (i) remain as a random dispersion in polymer-free fluid but (ii) aggregate side-by-side upon addition of PEO 100 kDa (0.1 wt%). Micrograph shows cells end-on, revealing the hexagonal symmetry of the side-by-side stacks.

One way to test this idea is to examine the reversibility of cable formation: given the entropic nature of the depletion attraction, we expect that after a cable has formed, removing polymer from the solution around it should allow the cable to disintegrate via random thermal motion of the cells. This is precisely what we observe upon flushing a suspension of cables with polymer-free liquid (Fig. 2b–c and Movie S7).

Support for the depletion attraction idea also comes from close inspection of the initial dynamics of cable formation. As exemplified in Movie S8 and Fig. S10 obtained using fast imaging, an initial “founder” cell grows and divides, giving rise to a pair of cells oriented end-to-end (Fig. S10a). Next, instead of thermally diffusing apart, the cells quickly lock in side-by-side (Fig. S10b). This preference for initially being arranged side-by-side versus end-to-end is directly predicted by the physics of the depletion interaction: the overlapping excluded volume *V*_ev_, and therefore the magnitude of the depletion potential *U*_dep_, are larger for a pair of cells when they are arranged side-by-side versus end-to-end [13, 75], as schematized in Fig. 2a(ii). Thus, in the absence of other constraints, over time scales shorter than the cellular doubling time *t*_d_, the polymer-induced depletion interaction causes pairs of cells to preferentially pack side-by-side. As a more direct test of this prediction, we culture cells in nutrient-free polymer solution, in which proliferation is arrested (*t*_d_ → ∞). Under these conditions, the cells indeed aggregate side-by-side, consistent with previous observations [9, 10, 12], forming hexagonally-ordered clusters (Fig. 2d). Taken altogether, these observations establish that depletion attraction induced by polymers plays a key role in cable formation. However, after an initial pair of cells has packed side-by-side, subsequent divisions typically lead to a persistent end-to-end configuration – thus resulting in the growth of cables. Why do these subsequent divisions not lead to side-by-side packing? We find the answer to this question using agent-based simulations.

### Agent-based simulations of a proliferating bacterial colony recapitulate cable formation in polymeric fluids

What biophysical features of proliferation in a polymer solution are essential for cable formation? We address this question using two-dimensional agent-based Brownian Dynamics simulations, detailed in the *Supplementary Information*. Each simulation begins with a dilute dispersion of cells, just as in the experiments, that proliferate in a polymer solution with a prescribed polymer size, viscosity, and osmotic pressure. The simulations incorporate four key features, schematized in Fig. 3a:

**Fig. 3:**
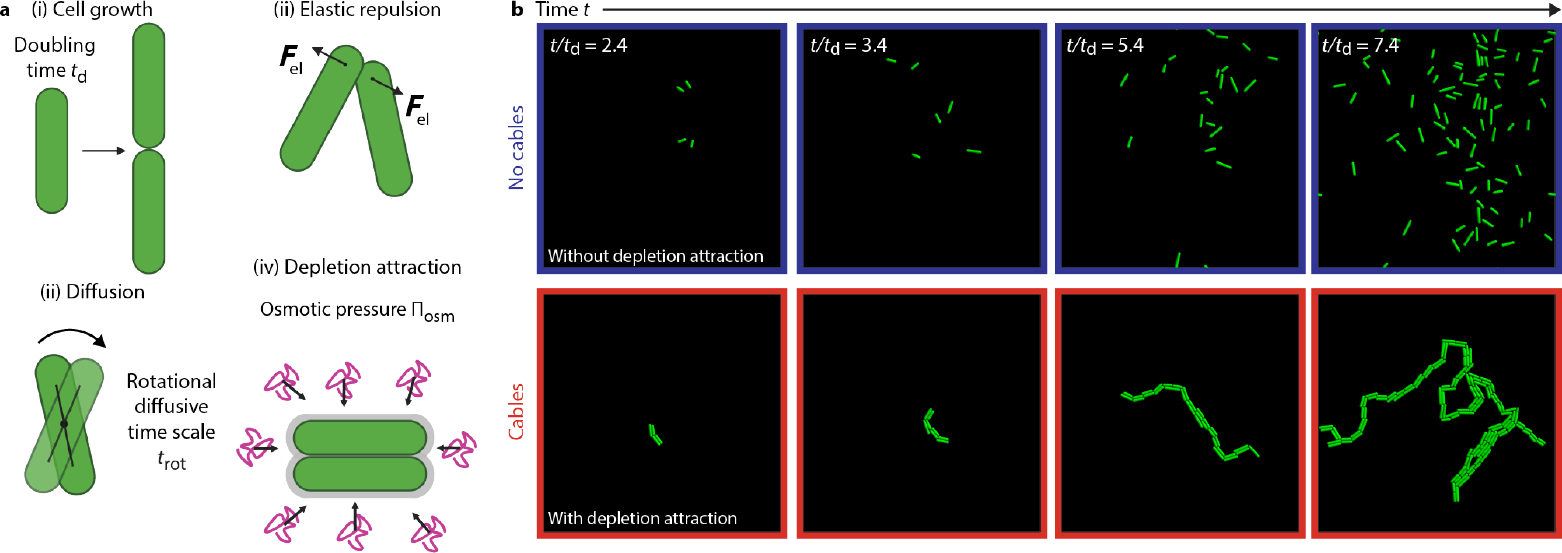
Agent-based simulations recapitulate cable formation in polymeric fluids. **a**, Schematic of the four key biophysical features incorporated into the agent-based simulations: (i) Cells are growing spherocylinders that exponentially elongate, divide, and separate, (ii) cells are stiff and elastically repel each other, (iii) cells are subject to translational and rotational diffusion due to thermal agitation, (iv) polymers induce a depletion attraction whose interaction potential depends on the geometry and relative orientation of adjacent cells. **b**, Time series of simulation results showing that in the polymer-free case, cells form a random dispersion, whereas with added polymer, proliferating cells form elongating cables, as in the experiments. Full dynamics can be seen in Movies S9–10.

1. The individual cells continually elongate exponentially, separating into two progeny after a doubling time *t*_d_.
2. Adjacent cells have a short-ranged elastic repulsion that pushes them apart and keeps them from overlapping.
3. Cells are subject to thermal agitation and undergo both translational and rotational diffusion in the viscous polymer solution, with corresponding diffusivities that are determined by the polymer solution viscosity and cell size/shape.
4. Cells are subject to the polymer-induced depletion attraction given by the pairwise interaction potential of magnitude *U*_dep_ = Π_osm_*V*_ev_, where Π_osm_ is a characteristic of the polymer solution to be tested and *V*_ev_ is set by the polymer size and cell sizes and orientations.

Each simulation can then be parameterized by two dimensionless quantities: *t*_rot_*/t*_d_, which compares the characteristic time scale for cells to rotationally diffuse to their doubling time, and *U*_ee_*/k*_B_*T*, which compares the magnitude of the depletion potential for cells arranged in the initial end-to-end configuration after doubling (Fig. 2a(ii)) to the characteristic thermal energy *k*_B_*T*, where *k*_B_ is Boltzmann’s constant and *T* is temperature.

Remarkably, these four key biophysical features are sufficient to recapitulate the experimental observations. An example is shown in Fig. 3b. In the absence of the depletion interaction (*U*_ee_ = 0 *k*_B_*T*, top row), cells continually grow, divide, separate, and thermally diffuse away, forming a random dispersion as in the experiments (Movie S9, Fig. 1a). By contrast, when the polymer-induced depletion interaction is incorporated (*U*_ee_ = 15.55 *k*_B_*T*), bottom row), the cells quickly lock in side-by-side and then continue to proliferate end-to-end, forming a large, multicellular cable just as in the experiments (Movie S10, Fig. 1b–e).

### Simulations reveal that the depletion interaction holds cells in a metastable end-to-end state after division, causing cables to form

Having demonstrated that the simulations can recapitulate cable formation, we next use them to address the question: Why exactly does the polymer-induced depletion attraction cause cells to proliferate in cables? Close inspection of the incipient stage of cable formation provides a clue: as shown in the initial frames of Fig. 3b (bottom) and Movie S10, a cable grows from an initial cluster of four cells, with two initially arranged in the energetically-preferred side-by-side configuration, but with their progeny retained end-to-end in an interdigitated configuration—as exemplified by the leftmost insets in Fig. 4a. Given that the most energetically-favorable configuration is for all cells to pack side-by-side [13, 75], we hypothesize that this interdigitated end-to-end configuration is a metastable state, enabled by the strength of the depletion interaction, that keeps the cells kinetically trapped in a cable.

**Fig. 4:**
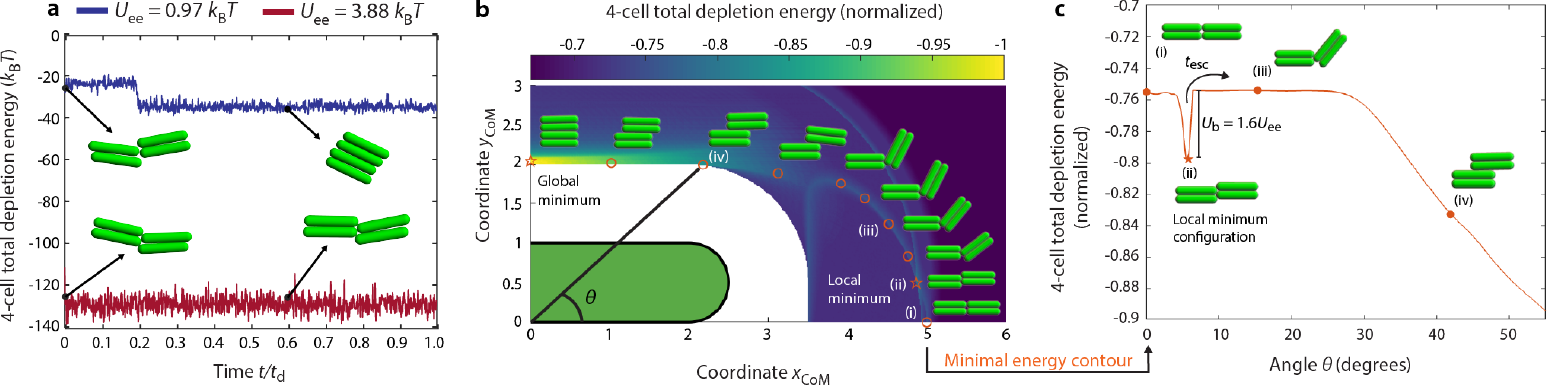
The depletion interaction holds cells in a metastable end-to-end state after division. Panels show the results of model and simulations (detailed in *Supplementary Information*) of four non-proliferating cells initially arranged in a proto-cable as labeled (i) in **b–c**. Cells quickly lock in to the interdigitated configuration labeled (ii). **a**, When the depletion attraction is strong (red), cells remain in this interdigitated configuration, whereas when the depletion attraction is weaker (blue), cells transition to the most energetically-preferred side-by-side configuration. Curves show the time series of the depletion potential, given by minus the product of the osmotic pressure and the overlapping excluded volume of a given configuration; the negative sign indicates an attractive interaction. **b**, Color map shows the landscape of the maximum possible depletion potential between 4 cells in different orientations for the case of *U*_ee_ = 0.97*k*_B_*T*. Two cells are fixed in the side-by-side configuration oriented along the *x*-axis, with their diameters spanning 0 < *y*_CoM_ < 1 and 1 < *y*_CoM_ < 2. The *x*_CoM_ and *y*_CoM_ coordinates describe the center of mass of the other two cells that can translate and rotate, again in the side-by-side configuration. The insets show ten distinct configurations at (*x*_CoM_, *y*_CoM_) positions indicated by the open symbols; stars indicate minima in the energy landscape. **c**, Minimal free energy contour from **b**, showing that the interdigitated end-to-end configuration (ii) is a metastable state characterized by a local free energy minimum given by *U*_b_ = 1.6*U*_ee_.

Simulations of four non-proliferating cells initially arranged both end-to-end and side-by-side, as shown by (i) in Fig. 4b–c, support this hypothesis. The cells quickly lock in to the interdigitated configuration shown by (ii) in Fig. 4b–c. Then, if the depletion interaction is strong (*U*_ee_ = 3.88 *k*_B_*T*, red curve in Fig. 4a), the cells remain trapped in this configuration over the entire duration of the doubling *t*_d_—ultimately enabling them to proliferate into a cable. However, when the depletion interaction is weaker (*U*_ee_ = 0.97 *k*_B_*T*), thermal diffusion allows the cells to eventually reconfigure into the most energetically-favorable side-by-side configuration before they double, as shown by the abrupt dip in the blue curve. These observations suggest that the interdigitated end-to-end configuration is indeed a metastable state.

Mapping the full orientation-dependent landscape of the potential due to the depletion interaction (magnitude *U*_dep_ = Π_osm_*V*_ev_), indicated by the colors in Fig. 4b, confirms this suggestion. As expected, having all four cells arranged side-by-side is the most energetically-favorable state, reflected by the global minimum in the free energy landscape (leftmost star). However, as we hypothesized, the interdigitated end-to-end configuration is a local minimum in this free energy landscape (rightmost star). Plotting the contour of minimal energy that extends across this landscape (Fig. 4c) reveals this local minimum, of depth ≈ 1.6*U*_ee_, even more clearly.

### A morphological state diagram unifies the interplay between depletion attraction and thermal diffusion in controlling cable formation

A clear picture of the biophysical rules underlying cable formation thereby emerges: After a side-by-side pair of cells divides end-to-end, shown by (i) in Fig. 4c, they remain kinetically trapped in the metastable interdigitated end-to-end configuration (ii) for an approximate escape time *t*_esc_, which is determined by cell size and shape and polymer size, concentration, osmotic pressure, and viscosity. If *t*_esc_ > *t*_d_, the cells continue to proliferate in this configuration, ultimately generating a cable. If *t*_esc_ < *t*_d_, however, thermal energy enables the cells to escape this kinetic trap and continue to explore other configurations, as shown by (iii)–(iv), and they do not form cables. This biophysical picture unifies all our experimental observations demonstrating that cable formation arises for diverse cell types, and across diverse polymer solutions when they are sufficiently concentrated, in a reversible manner.

As a final quantitative test of this picture, we directly compute *t*_esc_ for different polymer solutions following Kramers’ seminal work on thermally-activated escape from a potential well [76]. The boundary corresponding to *t*_esc_ = *t*_d_, which separates the two different regimes predicted by our analysis, is then given by the solid black curve in Fig. 5a. This plot represents a morphological state diagram spanned by the two dimensionless control parameters revealed by our analysis, *t*_rot_*/t*_d_ and *U*_ee_*/k*_B_*T*; to the right of the black boundary, we expect that cells proliferate into cables (red-outlined panels in Fig. 5b), whereas to the left of it, they do not (blue-outlined panels). We also expect a transitional regime immediately to the left of the black boundary, for which *t*_esc_ < *t*_d_ and cells are not kinetically trapped in the end-to-end configuration as they double, but *U*_ss_ ≳ *k*_B_*T*, causing cells to still stack side-by-side in small aggregates (light blue). As summarized by the triangles (experiments) and circles (simulations) in Fig. 5a, with representative colony morphologies shown in Fig. 5b, our data fully confirm these expectations. Thus, taken altogether, our experiments, simulations, and theoretical analysis establish biophysical principles that describe how bacterial colonies proliferate in polymeric solutions—across diverse cell types and polymers, both natural and synthetic.

**Fig. 5:**
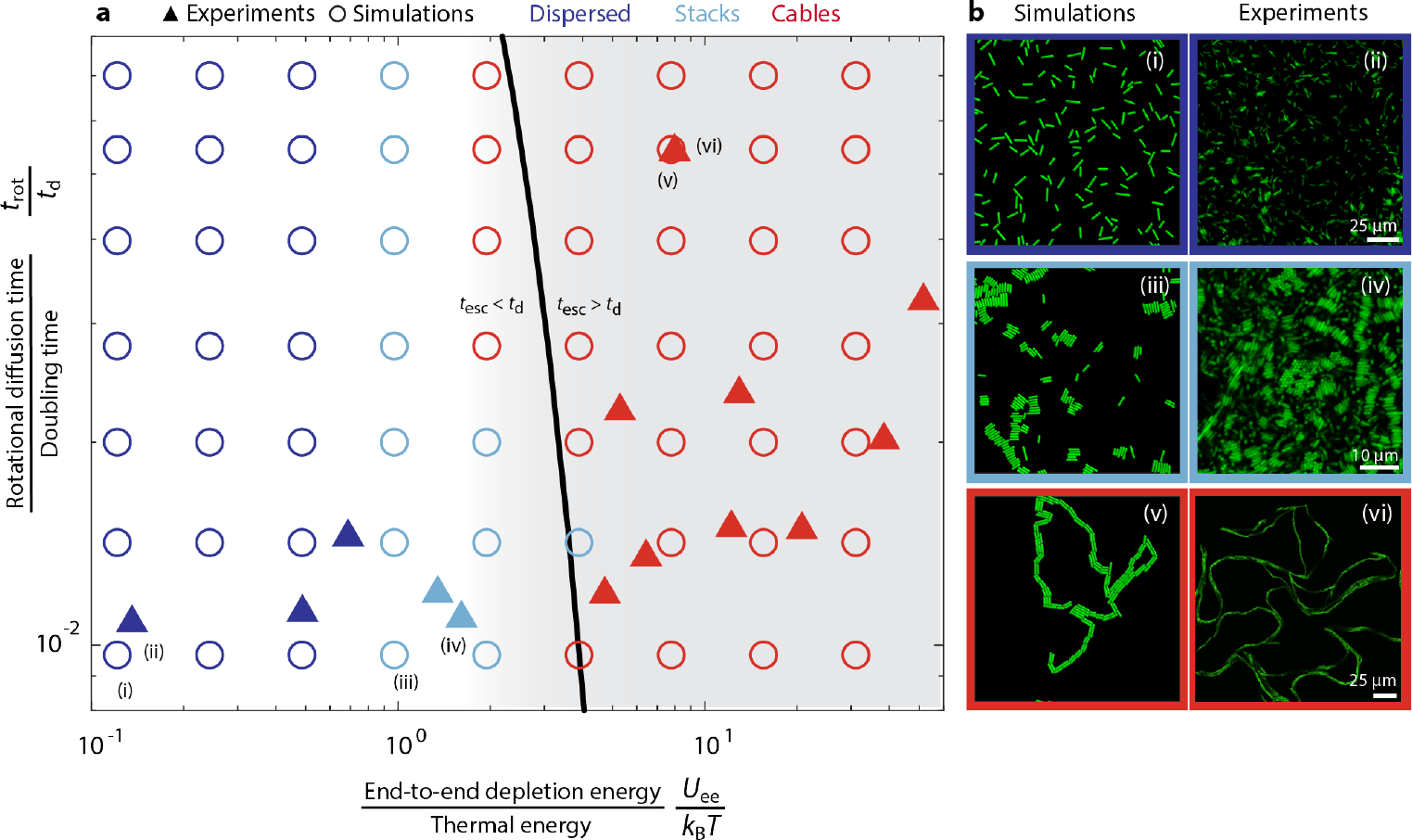
A morphological state diagram unifies the interplay between depletion attraction and thermal diffusion in controlling cable formation. **a**, State diagram is spanned by the two dimensionless parameters *t*_rot_*/t*_d_ and *U*_ee_*/k*_B_*T*, where the former compares the characteristic time scale for cells to rotationally diffuse to their doubling time, and the latter compares the magnitude of the depletion potential for cells arranged in the end-to-end configuration to the thermal energy; both are calculated as described in the *Supplementary Information*. Filled triangles and open circles show experimental and simulation results, respectively. Solid black curve shows our theoretical prediction for the onset of cable formation, given by *t*_esc_ = *t*_d_, where *t*_esc_ is the typical duration spent in the metastable interdigitated end-to-end configuration (given by Eq. S46) and *t*_d_ is the doubling time. Both agent-based simulation and *t*_esc_ shown in this figure assume a polymer size or polymer radius of 75 nm. This polymer radius corresponds, approximately, to the median size of polymer experimentally used in this study. The shaded grey region indicates our estimate for the region of this state diagram that describes in situ host mucus (detailed in *Supplementary Information*); interestingly, this region coincides with the onset of cable formation. Dark blue, light blue, and red symbols indicate colonies that formed random dispersions, small side-by-side stacks, and cables, respectively. **b**, Snapshots of representative colony morphologies for each of these three states. (i) *U*_ee_ = 0.25 *k*_B_*T, t*_rot_ = 0.009 *t*_d_; (ii) 0.01 wt% Ficoll 400 kDa; (iii) *U*_ee_ = 0.97 *k*_B_*T, t*_rot_ = 0.009 *t*_d_; (iv) 0.01 wt% NaCMC 250 kDa; (v) *U*_ee_ = 7.77 *k*_B_*T, t*_rot_ = 0.05 *t*_d_; (vi) 0.25 w/w% PEO 5 MDa.

## 3 Discussion

Despite their prevalence in natural habitats, little is known about how polymers influence one of the most fundamental aspects of bacterial life—their proliferation in colonies. By combining experiments, simulations, and theory, we have shown that extracellular polymers can shape bacterial colonies through purely physicochemical interactions, as broadly suggested by previous simulations [77]. In particular, we found that non-motile bacteria proliferating in sufficiently concentrated polymer solutions—both biological and synthetic—nematically align to form an intertwined network of long, serpentine, multicellular cables. This characteristic colony morphology arises due to the combined influence of polymer-induced entropic attraction holding cells together and the enhanced solution viscosity hindering cells from diffusively separating after dividing. Hence, this phenomenon arises generically and predictably across diverse bacterial species and polymer compositions; indeed, not only did our experiments directly demonstrate cable formation in gut mucus, but our analysis (detailed in *Supplementary Information*) indicates that, more broadly, biological polymers like mucus have physicochemical properties that promote cable formation, as shown by the grey region of Fig. 5. Our work thus uncovers new quantitative principles governing the morphogenesis of bacterial colonies—and potentially other microbial systems—in their complex environments in the real world. It also opens up a new direction for research in soft matter physics: while polymer-induced entropic attraction is well-studied for passive particulate systems, our work builds on previous simulations of biofilms [77] to highlight that fascinating new behaviors can emerge when the constituent particulates can also proliferate [15].

Because cables arise generically due to entropic forces induced by polymers on cells, they are fundamentally different from the “chains” or “cords” that sometimes arise for certain bacterial species, even in polymer-free conditions, due to specific biochemical interactions or cellular processes. One prominent example arises when host-secreted antibodies cross-link proliferating cells of *Salmonella enterica* together [78]. Another example is thought to arise for *Mycobacterium tuberculosis* due to hydrophobic interactions between mycomembrane lipids that strongly adhere cells together [79]. Similarly, under certain conditions, *E. coli* surface adhesins can adhere cells together in chains during biofilm formation [80]. In all three of these cases, specific biochemical interactions prevent cells from separating after division—unlike in cable formation, which arises due to a fundamentally different physicochemical mechanism across a broad range of polymer chemistries. A useful direction for future work will be to investigate how cable formation may be altered by the added influence of such specific biochemical interactions. A final example is the chains of cells that arise during *Bacillus subtilis* biofilm formation; unlike our cables, in which cells fully divide and separate from each other during proliferation, these chains form because cells do *not* completely divide and separate, and are therefore inseparably retained end-to-end [81–83]. Our work thus reveals a distinct, more general mechanism by which proliferating bacteria can form such long, multicellular cables when exposed to polymers.

Extensive work has focused on the ability of polymers to enhance the mobility of swimming bacteria [5–8]. By contrast, their influence on non-motile cells is understudied, despite the fact that many bacteria in natural polymeric environments (e.g., mucus) are non-motile or lose motility [33–56]—an important virulence factor that often correlates with pathogenesis and colonization/biofilm formation. Our study therefore focused on the proliferation of non-motile cells. However, a natural extension of our work is to investigate how cellular motility may alter cable formation. Intriguingly, cables are not formed when we repeat our experiments with swimming *E. coli* (Movies S11 and S12)—presumably because the hydrodynamic force generated by swimming cells exceeds the attractive depletion force induced by polymers, as supported by calculations in the *Supplementary Information* [12, 84]. Moreover, in the case of mucus, additional biochemical interactions can further promote dispersal of swimming cells [85]. Further investigating the interplay between cellular motility and polymer-induced cable formation will thus be an interesting direction for future research. Also, given that the entropic forces underlying cable formation depend sensitively on cell size and shape, another natural extension of our work is to investigate how cables form for cells of other shapes (e.g., curved) and in mixtures of different founder cell types.

What are the biological implications of cable formation? This phenomenon could be beneficial to non-motile bacteria by giving their colonies a way to extend outward and explore new environmental niches, including on surfaces or within host tissues [79, 86]. Proliferating in a cable could also help bacteria counter host immune responses against them, such as by impeding phagocytosis [87], compressing host cell structures [79], or protecting cells within a cable from antimicrobials [79]. Alternatively, it could be that hosts secrete polymers, such as mucins, to force cells to proliferate in cables, potentially enhancing clonal extinction rates and pathogen clearance [78], in addition to specific biochemical interactions that downregulate pathogenic genes [1, 59, 88]. Such effects may also be induced by dietary polymers transiting through the gut [89, 90], potentially providing an indirect link between host diet, the physiology of the gut microbiome, and host health. Moreover, by localizing cells together, cable formation could alter the dynamics and extent of infection by bacteriophages throughout a colony, as suggested by simulations [91]. Building on our findings to investigate these possible consequences of cable formation will be an exciting direction for future research.

## 4 Methods

### 4.1 Preparation of cells

#### E. coli

We incubate an overnight culture of *E. coli* strain W3110 in 2 wt% Lennox Lysogeny Broth (LB) at 30°C. Next, we inoculate 20 *µ*L from the overnight culture in 2 mL of 2 wt% LB for 3 hours such that, at the time of imaging, bacteria are in mid–exponential phase. We use a strain with a deletion of the flagellar regulatory gene *flhDC* that renders the cells non-motile; we verify these cells are non-motile using direct visualization.

#### P. aeruginosa

We incubate an overnight culture of *P. aeruginosa* strain PA01 in 2 wt% LB, supplemented with 200 *µ*g/mL of Carbenicilin to preserve the strain’s fluorescence, at 37°C. Next, we inoculate 20 *µ*L from the overnight culture in 2 mL of 2 wt% LB for 3 hours. We use a strain with a double *fliC* and *pilA* deletion that renders the cells non-motile; we verify these cells are non-motile using direct visualization.

#### V. cholerae

We incubate an overnight culture of *V. cholerae* O1 biovar El Tor strain C6706 in 2 wt% LB at 37°C. Next, we inoculate 20 *µ*L from the overnight culture in 2 mL of 2 wt% LB for 5 hours with the addition of rolling beads in culture to keep the bacteria dispersed. We use a strain with several gene deletions: It has a deletion of *pomA*, which renders the cells non-motile; we verify these cells are non-motile using direct visualization. It also has deletions of *rbmA, bap1, rbmC*, and *vpsL*, which renders the cells as non-biofilm formers.

### 4.2 Preparation of polymer solutions

#### Synthetic polymers

We obtain PEO 5 MDa from ColorCon and Sigma Aldrich; PEO 1 MDa and 100 kDa from ColorCon; Ficoll 400 kDa from Research Products International; and Carboxymethyl Cellulose Sodium Salt from MP biomedicals. We prepare polymer stock solutions in 2 wt% LB and then put them on a spinning rotor for at least 8 hours until the solution is optically clear. The PEO solutions are additionally filtered using a 5 *µ*m-pore size membrane from GE Healthcare life sciences. We use dilutions of these concentrated stock solutions in LB to prepare polymer solutions at defined concentrations.

#### Human colonic mucus

We use mucus from human primary transverse colon cells [57] obtained from Altis biosystems. In particular, the mucus is siphoned from the surface of a culture of resident goblet cells (Altis RepliGut Planar transverse colon model), and then directly frozen and stored at -20°C. We then defrost and mix this mucus stock solution with 10 w/w% LB at an appropriate ratio such that the final LB concentration is 2 w/w% and the final mucus concentration is 0.5 w/w%.

#### Muc2 mucins

We use purified native porcine gel-forming intestinal mucin (Muc2) [1, 2] obtained from the Ribbeck lab at MIT as a lyophilized powder. This powder is dissolved in autoclaved distilled water at a stock concentration of 1 w/w%. We then leave this solution on a spinning rotor at 4°C overnight a day prior to using it in an experiment. The day of the experiment, we mix the Muc2 stock solution with 10 w/w% LB and use dilutions with autoclaved distilled water such that the final LB concentration is 2 w/w% and a the final Muc2 concentration is 0.5 w/w%. We then leave the test solution on a spinning rotor for at least 2 hours prior to use in the experiment to ensure a homogeneous solution.

### 4.3 Imaging bacterial proliferation

#### *E. coli* in synthetic polymer solutions

We fill the base (20 mm diameter, 1 mm height) of a transparent-walled glass-bottom Petri dish, 35 mm in diameter and 10 mm in height overall, with a test solution containing cells at an initial concentration of ≈ 4 × 10^5^ cells/mL and polymer at the concentration to be tested. Next, we seal this base with an overlying circular Polydimethylsiloxane (PDMS) slab to minimize evaporation while still allowing oxygen to be available. We then image the sealed chamber from below using a Nikon A1R+ inverted laser scanning confocal microscope with the stage maintained at 30 ± 1°C. We acquire fluorescent optical slices throughout the depth of the sample every 1 to 5 min over a total duration of ≈ 6 to 14 hours.

For the experiment in Fig. 2b–c investigating the reversibility of cable formation, we again use a transparent-walled glass-bottom Petri dish sealed with an overlying PDMS slab as described above. However, we additionally pierce the PDMS slab with three 20 in gauge needles, sealed using optical glue, and connected to a Harvard Apparatus 11 Elite syringe pump to provide fluid in/outflow. In particular, one needle acts as an inlet for injection of the test solution containing ≈ 4 × 10^5^ cells/mL suspended in 0.1 w/w% PEO 100 kDa as the polymer; the second acts as an inlet for injection of polymer-free LB media; and the third acts as the outlet. We then image the sealed chamber from below using a Nikon A1R+ inverted laser scanning confocal microscope with the stage maintained at 30 ± 1°C as before, first under quiescent no-flow conditions with the chamber filled with the test solution containing cells and polymer. Once cables have formed, we then pump polymer-free fluid through at a flow rate of 5 *µ*L/min to remove polymer-containing solution. Our experiments shown in Movie S13 (snapshots in Fig. S11) indicate that these flow rates are insufficient to fragment cables due to hydrodynamic forces.

For the experiment in Fig. 2d using non-proliferating cells, we inoculate 20 *µ*L from an overnight culture of cells into 2 mL of 2 wt% LB and let the bacteria grow for 3 hours in a shaking incubator until they reach mid-exponential phase. We then mix 10 *µ*L of the culture with a solution of PEO 100 kDa dissolved in 1 × Difco M9 Minimal Salts without a nutrient source at a polymer concentration of 0.1 w/w% and a cellular concentration of ≈ 6 × 10^7^ cells/mL. We then again fill the base of a transparent-walled glass-bottom Petri dish with the solution, sealed with an overlying PDMS slab as described above, and incubate the dish at 30°C in a static incubator for 2 hours. We then image the sealed chamber from below using a Nikon A1R+ inverted laser scanning confocal microscope with the stage maintained at 30 ± 1°C.

#### *E. coli* in human colonic mucus

We fill a custom-made PDMS rectangular channel (22 mm long, 2 mm wide, 25 or 90 *µ*m high) with the test mucus solution containing cells at an initial concentration of ∼3 × 10^6^ cells/mL. We then image the sealed channel from below using a Nikon A1R+ inverted laser scanning confocal microscope with the stage maintained at 33 ± 1°C. We acquire fluorescent optical slices throughout the depth of the sample every 3 min over a total duration of ≈ 12 hours.

#### *E. coli* in Muc2 solutions

We fill a glass capillary (4 cm long, 5 mm wide, 500 *µ*m high) with the test Muc2 solution containing cells at an initial concentration of ∼ 3 × 10^5^ cells/mL. We then image the capillary, sealed on both ends with paraffin oil to minimize evaporation while still allowing oxygen to be available, from below using a Nikon A1R+ inverted laser scanning confocal microscope with the stage maintained at 30 ± 1°C. We acquire fluorescent optical slices throughout the depth of the sample every 3 min over a total duration of ≈ 12 hours.

#### *P. aeruginosa* and *V. cholerae* in polymer solutions

We fill individual wells of a glass-bottom 96-well plate with 200 *µ*L of a test polymer solution and 0.2 *µ*L of the inoculum of cells such that their initial concentration is ~ 3 to 6 × 10^6^ cells/mL. We store the plate in a static 37°C incubator and image the plate at different time points: For *P. aeruginosa* we image at 210 min and 420 min after the start of incubation and for *V. cholerae* at 135 min and 303 min after incubation as shown in Fig. S8. For all images taken we use a 20× air objective mounted in a Nikon A1R+ inverted laser scanning confocal microscope with the stage maintained at 37 ± 1°C.

## Acknowledgments

N.S.W. acknowledges support from NSF Center for the Physics of Biological Function Grant PHY-1734030 and NIH Grant R01 GM082938. S.S.D. acknowledges support from NSF Grants CBET-1941716, DMR-2011750, and EF-2124863 as well as the Camille Dreyfus Teacher-Scholar and Pew Biomedical Scholars Programs, the Eric and Wendy Schmidt Transformative Technology Fund, and the Princeton Catalysis Initiative. We thank Tapomoy Bhattacharjee for assistance with a preliminary version of the experiments at the inception of this project and Brianna Royer for assistance with imaging. We also thank Tapomoy Bhattacharjee, Scott Chimileski, Zemer Gitai, Sridhar Mani, and Jessica Mark Welch for useful discussions; Hao Li, Sridhar Mani, and Altis Biosystems for providing mucins obtained from human primary transverse colon cells; the laboratory of Howard Stone for providing access to the rheometer; and the laboratories of Bob Austin, Bonnie Bassler, and Zemer Gitai for providing strains of *E. coli, V. cholerae*, and *P. aeruginosa*, respectively.

## Author contributions

S.G.L.C. performed all experiments, theoretical calculations, agent-based simulations, and analyses; C.A.S. and G.C-O prepared and contributed mucins and assisted with experimental design; K.R. contributed mucins and assisted with the interpretation of results; N.S.W. assisted in research design and analysis/interpretation of the results; S.S.D. conceived/designed the overall study and assisted in research design and analysis/interpretation of the results; S.G.L.C., K.R., N.S.W., and S.S.D. contributed to writing the paper.

## Competing interests

The authors declare no competing interests.

## Supplementary movies

All movies are available at https://zenodo.org/doi/10.5281/zenodo.10966670.

1. **Movie S1:** Non-motile *E. coli* proliferating in polymer-free LB media.
2. **Movie S2:** Non-motile *E. coli* proliferating in LB media supplemented with 0.5 w/v% mucus.
3. **Movie S3:** Non-motile *E. coli* proliferating in LB media supplemented with 0.5 w/w% Muc2 mucin polymer.
4. **Movie S4:** Non-motile *E. coli* proliferating in LB media supplemented with 1 w/w% PEO 100 kDa.
5. **Movie S5:** Non-motile *E. coli* proliferating in LB media supplemented with 5 w/w% Ficoll 400 kDa.
6. **Movie S6:** Non-motile *E. coli* proliferating in LB media supplemented with 1.0 w/w% NaCMC 250 kDa.
7. **Movie S7:** Non-motile *E. coli* proliferating in LB media supplemented with 0.1 w/w% PEO 100 kDa. At the start of the video (*t* = 0 min) we flush a suspension of cables with polymer free liquid at a flow rate of 5 *µ*L/min, thus decreasing the polymer concentration as time increases.
8. **Movie S8:** Non-motile *E. coli* proliferating in LB media supplemented with 0.2 w/w% PEO 5 MDa.
9. **Movie S9:** Agent-based simulation of non-motile *E. coli* cells proliferating without depletion attraction. Parameters: *t*_rot_ = 0.05 *t*_d_ and *U*_ee_ = 0 *k*_B_*T*.
10. **Movie S10:** Agent-based simulation of non-motile *E. coli* cells proliferating with depletion attraction. Parameters: *t*_rot_ = 0.05 *t*_d_ and *U*_ee_ = 15.55 *k*_B_*T*.
11. **Movie S11:** Maximum intensity projections of swimming *E. coli* cells proliferating in polymer-free LB media.
12. **Movie S12:** Maximum intensity projections of swimming *E. coli* cells proliferating in LB media supplemented with 0.5 w/w% PEO 5 MDa.
13. **Movie S13:** Non-motile *E. coli* proliferating in LB media supplemented with 0.1 w/w% PEO 100 kDa. Twenty minutes into the video (*t* = 20 min) we flush a suspension of cables with 0.1 w/w% PEO 100 kDa polymer solution at a flow rate of 5 *µ*L/min keeping the polymer concentration constant as time increases.

## Supplementary information

### Characterization of polymer solutions

We use an Anton Paar MCR501 rheometer with both cone-plate and double-gap geometries to characterize our polymer solutions. In particular, we measure the dynamic shear viscosity as a function of imposed shear rate; as shown in Fig. S1, the viscosity does not vary appreciably with shear rate. Hence, we use the mean of the viscosity values measured at different shear rates to define the zero shear viscosity *η*.

**Fig. S1:**
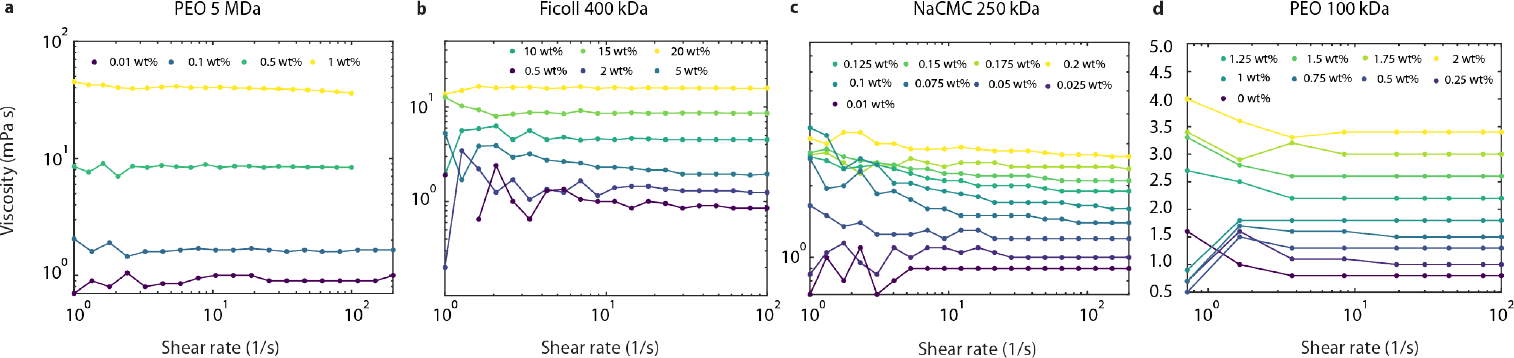
Flow curves of polymer solutions. **a-d**, The panels show the viscosity of the following polymer solutions: **a** PEO 5MDa, **b** Ficoll 400 kDa, **c** NaCMC 250 kDa, and **d** PEO 100 kDa as a function of shear rate at different polymer concentrations within the range of 0.72 − 200 s^*−*1^.

Measuring the dependence of this viscosity on polymer mass concentration *c* yields a straightforward way to determine the polymer overlap concentration *c*^∗^. In particular, writing the viscosity as a virial expansion *η* = *η*_s_ 1 + [*η*] *c* + *k*_H_ [*η*]^2^ *c*^2^ + … and retaining only the first three terms on the right hand side yields two distinct relations [73]; here, [*η*] is known as the intrinsic viscosity, *k*_H_ is a constant known as the Huggins coefficient, and *η*_s_ is the viscosity of the solvent (LB). The first, known as the Huggins equation, directly follows from rearranging this equation:

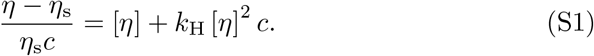

The second, known as the Kraemer equation, follows from the approximation that ln 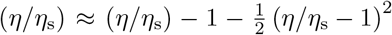; substituting from the virial expansion for *η/η*_s_ then yields the Kraemer equation:

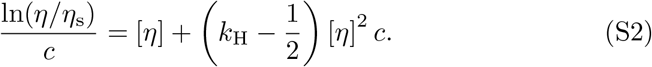

Fitting both equations to our data and extrapolating to *c* = 0 (Fig. S2) then yields a direct determination of the intrinsic viscosity; we then use the established relation *c*^∗^ = 0.77*/* [*η*] [92] to directly compute *c*^∗^ for each test solution, as given in Table S1.

**Fig. S2:**
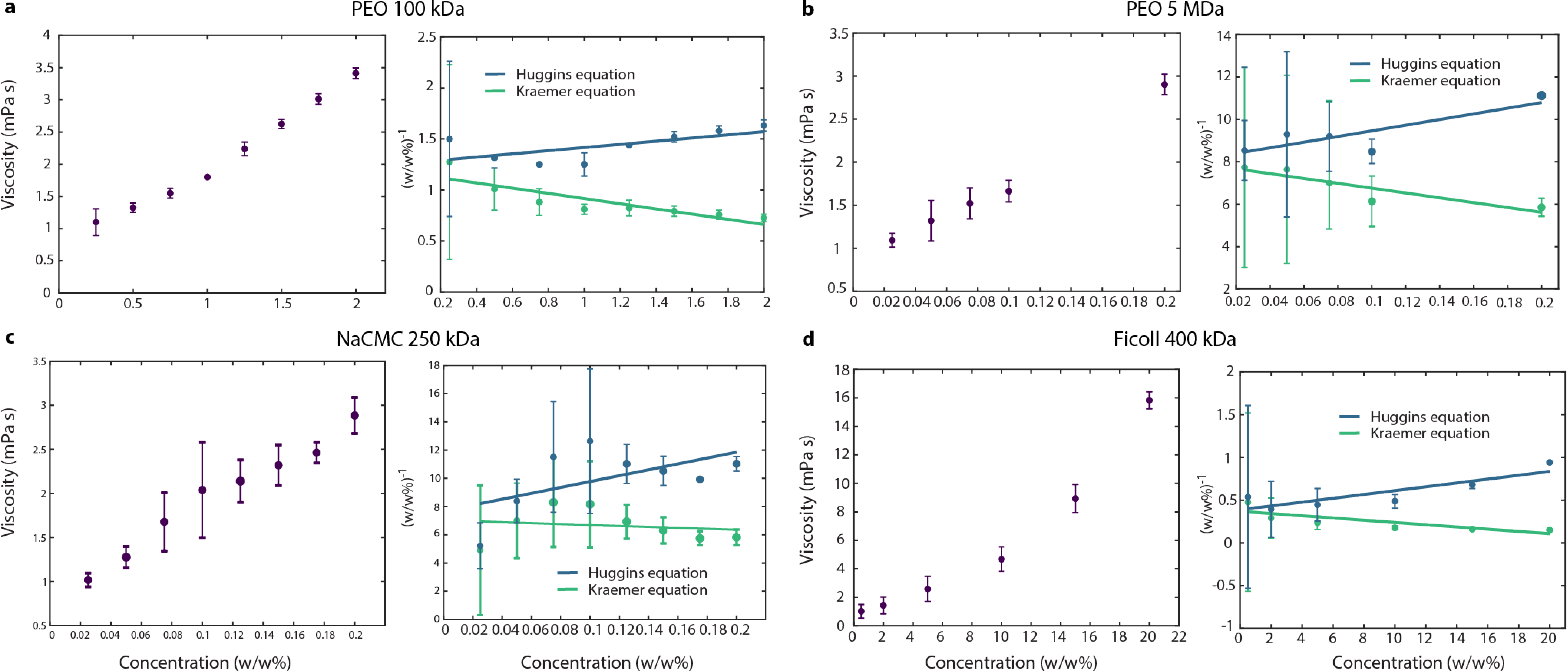
Overlap concentration measurements of polymer solutions. **a-d**, The panels show both the low shear rate viscosity vs. concentration curve for each polymer as well as its corresponding Huggins and Kraemer representations (dark blue and light green plots, respectively). Shear rheology showed that the viscosity of these dilute polymer solutions does not depend appreciably on shear rate, hence the dots represent the mean viscosity measured over all shear rates at constant polymer concentration, while the error bars correspond to the variation of the viscosity at constant polymer concentration over the range of shear rates used, 1−200 Hz. We used the Huggins and Kraemer representations because we extrapolated the value of these curves to *c* = 0 in order to acquire the polymer solutions’ intrinsic viscosity, [*η*]. After extrapolating [*η*] we estimated an overlap concentration via the relation *c*^∗^ = 0.77*/*[*η*].

Having determined the overlap concentration, *c*^∗^, we next determine two other key parameters of each test polymer solution: the osmotic pressure Π_osm_ and the concentration-dependent polymer radius *R*_p_, both also given in Table S1. The former is given directly by the established relation [73]

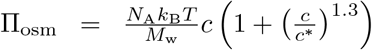, where *N*_*A*_ is Avogadro’s number, *k*_*B*_ is Boltzmann’s constant, *T* is temperature, and *M*_*w*_ is the polymer molecular weight. The latter is then given by the established relation [93,94] 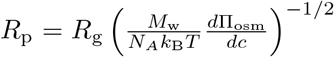, where 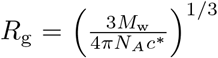 is the dilute polymer radius of gyration [73].

**Table S1:** Summary of experiments with *E. coli*. Quantities listed are: polymer concentration (*c*), overlap concentration (*c*^∗^), viscosity (*η*), osmotic pressure (Π_osm_), doubling time (*t*_d_), and polymer radius (*R*_*p*_). We also note whether cables were experimentally observed. The doubling time indicated by ^†^ was obtained by linearly interpolating the doubling times measured as a function of polymer concentration.

| Polymer | $c$<br>(w/w%) | $c^*$<br>(w/w%) | $\eta$<br>(mPa · s) | $\Pi_{\text{osm}}$<br>(Pa) | $t_d$<br>(min) | $R_p$<br>(nm) | Cables? |
| --- | --- | --- | --- | --- | --- | --- | --- |
| PEO 5 MDa | 0.01 | $0.09 \pm 0.01$ | $1.1 \pm 0.1$ | 0.05 | $35 \pm 1$ | 120 | No |
| PEO 5 MDa | 0.1 | $0.09 \pm 0.01$ | $1.6 \pm 0.4$ | 1.4 | $34 \pm 1$ | 67 | Yes |
| PEO 5 MDa | 0.25 | $0.09 \pm 0.01$ | $4 \pm 1$ | 5.4 | $36^\dagger$ | 43 | Yes |
| PEO 5 MDa | 0.5 | $0.09 \pm 0.01$ | $8.4 \pm 0.4$ | 23.4 | $39 \pm 1$ | 28 | Yes |
| PEO 5 MDa | 1 | $0.09 \pm 0.01$ | $40 \pm 2$ | 107.8 | $43 \pm 1$ | 18 | Yes |
| PEO 1 MDa | 1 | | $37 \pm 1$ | | | | Yes |
| PEO 100 kDa | 0.01 | $0.98 \pm 0.03$ | $0.90 \pm 0.03$ | 2.5 | $38 \pm 2$ | 16 | No |
| PEO 100 kDa | 0.1 | $0.98 \pm 0.03$ | $0.97 \pm 0.04$ | 27.2 | $37 \pm 3$ | 15 | Yes |
| PEO 100 kDa | 0.5 | $0.98 \pm 0.03$ | $1.3 \pm 0.1$ | 209.9 | $37 \pm 3$ | 11 | Yes |
| PEO 100 kDa | 1 | $0.98 \pm 0.03$ | $1.8 \pm 0.3$ | 668.5 | $40 \pm 2$ | 7 | Yes |
| Ficoll 400 kDa | 0.01 | $3.3 \pm 0.3$ | $0.90 \pm 0.03$ | 0.6 | $37 \pm 2$ | 17 | No |
| Ficoll 400 kDa | 0.1 | $3.3 \pm 0.3$ | $0.92 \pm 0.04$ | 6.4 | $36 \pm 3$ | 17 | No |
| Ficoll 400 kDa | 0.5 | $3.3 \pm 0.3$ | $1.0 \pm 0.1$ | 35.7 | $36 \pm 1$ | 14 | Yes |
| Ficoll 400 kDa | 1 | $3.3 \pm 0.3$ | $1.1 \pm 0.5$ | 83.8 | $37 \pm 3$ | 8 | Yes |
| Ficoll 400 kDa | 5 | $3.3 \pm 0.3$ | $2.6 \pm 0.9$ | 1176.5 | $40 \pm 2$ | 5 | Yes |
| NaCMC 250 kDa | 0.01 | $0.10 \pm 0.02$ | $0.9 \pm 0.1$ | 1.1 | $38 \pm 1$ | 43 | No |
| NaCMC 250 kDa | 0.1 | $0.10 \pm 0.02$ | $2.0 \pm 0.5$ | 25.4 | $39 \pm 2$ | 26 | Yes |
| Mucus | 0.5 |  |  |  |  |  | Yes |
| Muc2 | 0.5 |  |  |  |  |  | Yes |

### Identification of experimental cables using image analysis

To identify cables from the experimental confocal micrographs for Fig. 5, we:

1. Smooth the images using a 2D Gaussian filter.
2. Binarize the images using Otsu’s method of thresholding [95].
3. Skeletonize the largest object identified in the binarized images using the medial axis transform and thereby determine its backbone.
4. Identify a structure as a cable if the length of the backbone thereby identified exceeds 300 *µ*m, approximately 100 times the length of a cell body.

### Agent-based simulations

We model each cell *i* as an exponentially-growing spherocylinder comprised of a cylinder of length *l*_*i*_(*t*) and diameter *d*, capped on each’ end by a hemisphere of the same diameter:

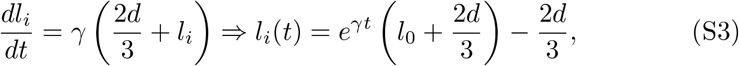

where *t* represents time elapsed after division, *l*_0_ is the initial cylinder length, and *γ* is the proliferation rate. Once the cell grows to twice its original length after a doubling time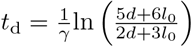, it is instantaneously replaced by two identical daughter cells; replacing the mother cell with two daughter cells in this manner results in a slight loss of total cell volume, but importantly, it does not increase overlap with any neighboring cells.

#### Equations of motion

To model the motion of each cell *i* we assume that each cell is subject to an elastic cell-cell repulsive force, a polymer-induced depletion attraction force, and a Langevin force to reflect thermal agitation.

The equations of motion, adopting Einstein notation, are given by:

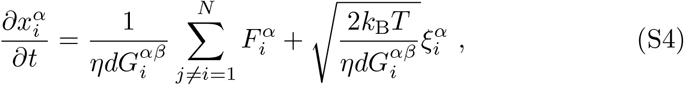

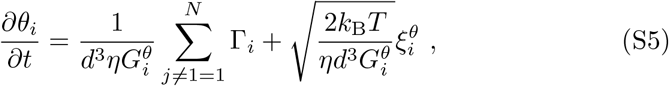

where *α* and *β* are indices representing the two axes that span the two-dimensional Euclidean space (*x, y*). The right hand side of Eqs. S4 and S5 have two terms: the first term accounts for cell-cell repulsive and polymer-induced forces and torques, while the second term represents the Langevin forces and torques. The variables 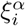 and 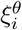 constitute random variables that obey ⟨*ξ*(*t*) ⟩ = 0 and ⟨*ξ*(*t*)*ξ*(*t*^*′*^) ⟩ = *δ*(*t − t*^*′*^) and have dimensions of inverse time to maintain dimensional consistency. In Eq. S4, 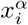 is a position coordinate of cell *i* and 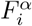 is a component of the net force on cell *i*, while in Eq. S5, *θ*_*i*_ is the orientation of cell *i* and Γ_*i*_ is the net torque on cell *i*. The coefficients 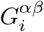 and 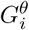 are cell *i*’s geometrical factors and have the following forms in the cell’s reference frame [96, 97]:

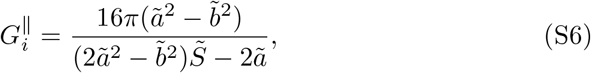

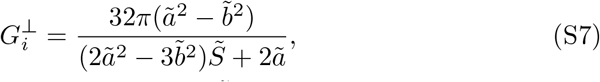

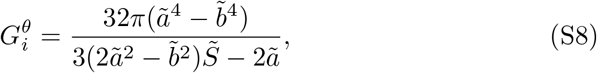

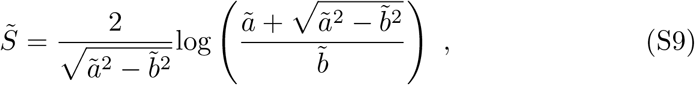

where Eqs. S6 and S7 correspond to geometrical factors that are parallel and perpendicular to the cell’s long axis, and Eq. S8 is the rotational factor. In the above equations we define *ã* (*l* + *d*)*/*2*d* and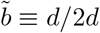.

We next express the forces in Eqs. S4 and S5 as the negative gradients of the relevant interaction potentials:

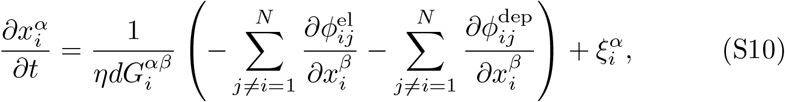

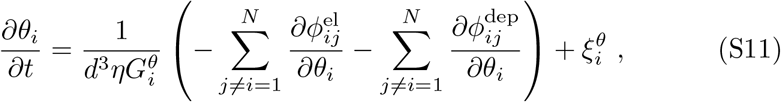

where 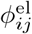 and 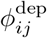 are the pairwise repulsive elastic and depletion attraction potentials between cell *i* and *j* respectively.

The elastic potential has the form [20, 98, 99]

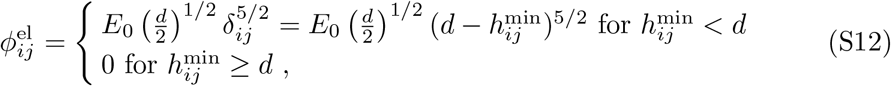

where *E*_0_ is the cell’s elastic modulus and *δ*_*ij*_ is the overlap distance of spherocylinders *i* and *j*, defined as the the shortest distance between the centerlines of spherocylinders *i* and *j* minus the diameter of the cells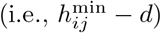.

The depletion attraction is described by the pairwise interaction potential

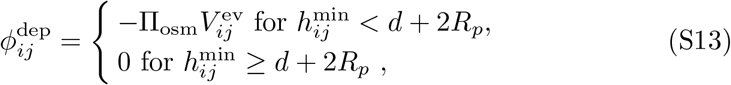

where Π_osm_ is the osmotic pressure of the polymer solution, *R*_*p*_ is the polymer radius, and 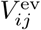 the overlapping excluded volume between two spherocylinders, given by [74]

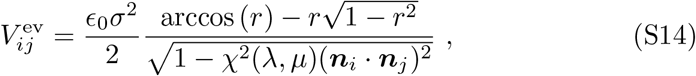

where *σ* = *d* + 2*R*_*p*_, 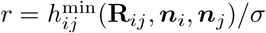 is the minimum distance between the centerlines of spherocylinders *i* and *j* divided by *σ*, ϵ_0_ is a fitting parameter, and *χ*^2^(*λ, µ*) is a function that has the form

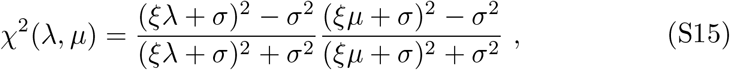

where in Eq. S15 the variable *ξ* is another empirical fitting parameter, while *λ* and *µ* are interaction lengths whose precise definitions are detailed in a previous study [74].

Therefore, Eqs. S10 and S11 can be written as

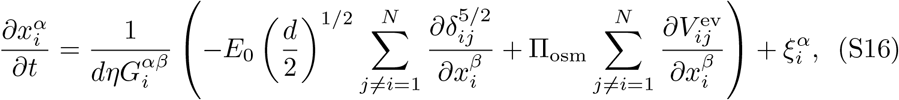

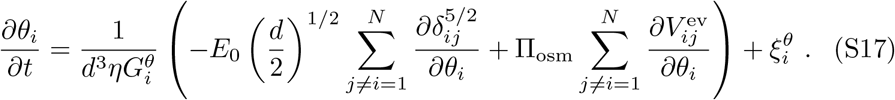

To nondimensionalize our equations we rescale the variables using characteristic time, length, and noise scales *t*_c_, *x*_c_, and *ξ*_c_:

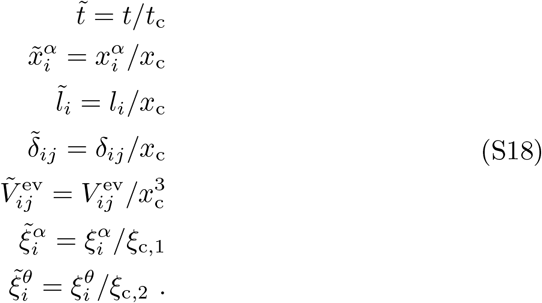

Substituting the above definitions into Eqs. S3, S10, and S11 yields:

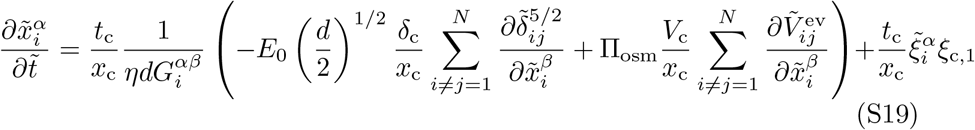

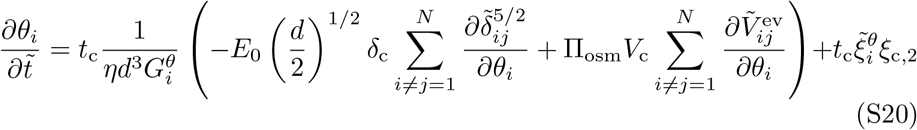

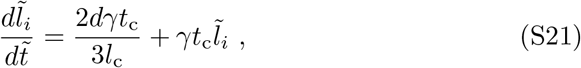

thereby defining the characteristic scales

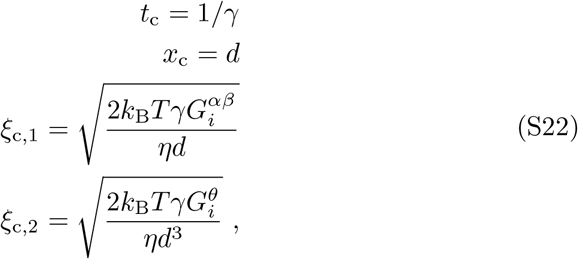

where 1*/γ* represents the characteristic proliferation time scale, *d* is the cell diameter, and the expressions for *ξ*_c,1_ and *ξ*_c,2_ stem from applying the fluctuation-dissipation relation for the diffusivity of a single cell. Hence, we arrive at the following dimensionless governing equations:

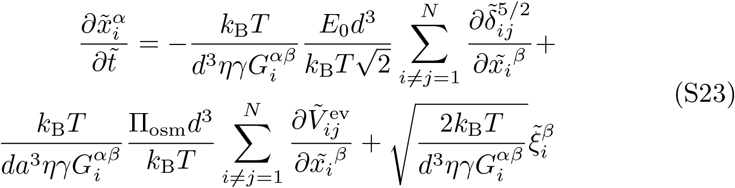

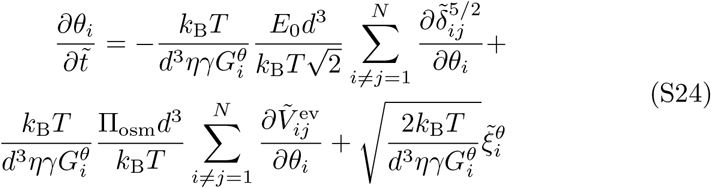

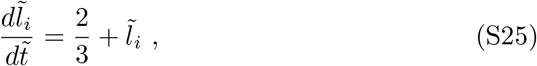

yielding the three dimensionless parameters

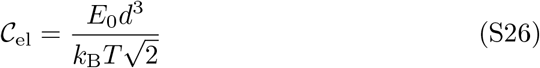

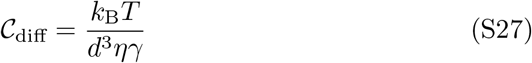

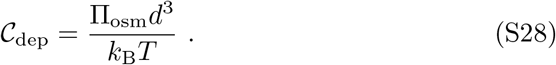

#### Physical significance of dimensionless parameters

As just described, nondimensionalization of the governing equations of motion for the agent-based simulations yields three dimensionless parameters describing this system. Below, we provide physical intuition for each.

The first, 𝒞_el_, can be rewritten as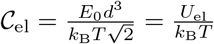, where the character-istic elastic energy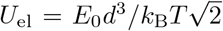. Hence, this parameter compares the characteristic elastic energy to thermal energy, and thus describes how easily a cell can be deformed by thermal fluctuations.

The second, 𝒞_diff_, can be re-written as 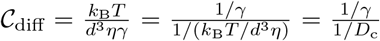.

Here, the numerator is the characteristic proliferation time, whereas the denominator is a characteristic rotational diffusive time scale. For the denominator to account for the anisotropic shape of a bacterium we multiply it by the rotational geometric factor for an ellipsoid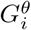; i.e., the rotational diffusive time scale is given by

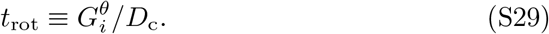

Hence, 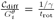 describes how rapidly a cell grows and divides compared to how rapidly it reorients due to thermal agitation. The inverse of this dimensionless parameter is the vertical axis of Fig. 5.

The third, 𝒞_dep_, can be re-written as 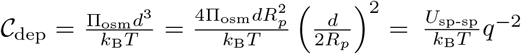, where 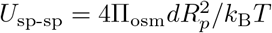 is the characteristic depletion energy associated with two spheres in contact and *q* = *d/*2*R*_*p*_ is the cell-polymer size ratio. Hence, this parameter describes the relative strength of depletion interactions compared to thermal fluctuations.

We also define a related dimensionless parameter when accounting for the finite elasticity of the cell surfaces in calculating depletion interactions between cells. In particular, we calculate the equilibrium length, *r*_eq_, between the centroids of the spheres of diameter *d* at which the sum of the depletion attraction energy, *U*_dep_, and the elastic repulsive energy, *U*_el_ are a minimum. The depletion energy and elastic energy have the form [90, 98]

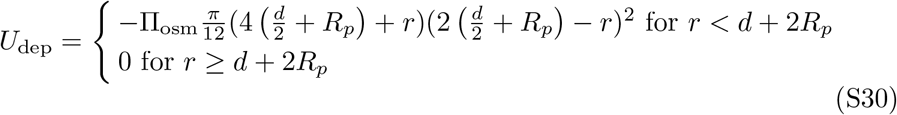

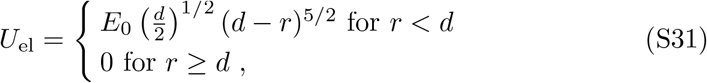

where *r* is the separation between the centroids of the two spheres and we define the total energy as *U*_tot_ = *U*_el_ + *U*_dep_. In the simulations, we numerically compute *U*_tot_ as a function of *r* to locate the equilibrium length, *r*_eq_, and extract the corresponding depletion energy, *U*_dep_(*r* = *r*_eq_) = *U*_ee_, to determine the dimensionless parameter shown in Fig. 5.

#### Numerical implementation of agent-based simulations

We use a custom C++ script where we implement an explicit Euler-Maruyama method to numerically solve the nondimensional equations derived previously. Discretizing S23, S24, and S25 yields

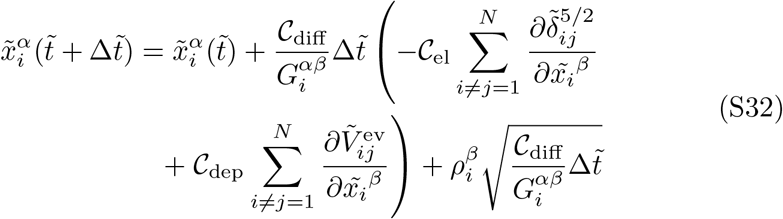

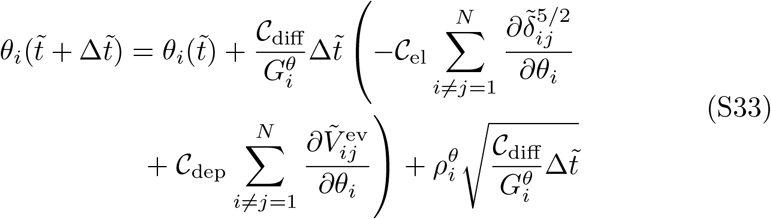

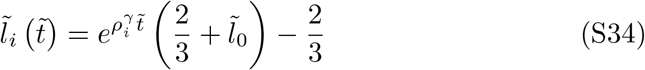

where 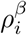 and 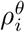 are random numbers selected from a Gaussian distribution of mean zero and variance one, 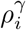 is a random number selected from a Gaussian distribution of mean 1 and standard deviation 0.2, and 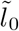 is the cell cylinder length at 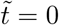.

To evolve the equations of motion and proliferation in time we implement the following protocol:

1. At each time step, elongate and divide cells following equation S34.
2. Compute the net force, 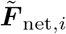, and net torque, Γ_net,*i*_, on each cell *i* given the potentials in equations S12 and S13.
3. Transform the net force on cell *i* to each cell *i*’s own reference frame by performing dots products between the net force vector, 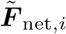, and unit vectors corresponding to a cell’s orientation vector, ***n***_*i*_, and an in-plane vector, ***o***_*i*_, that is orthogonal to ***n***_*i*_

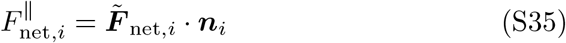

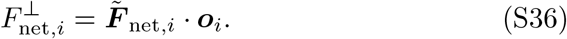
4. Compute the translational displacement of cell *i* in its own reference frame

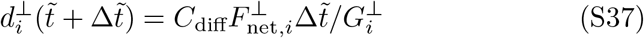

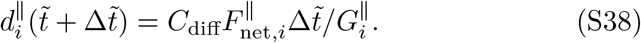
5. Transform displacements into the lab frame to update cell *i*’s position accordingly

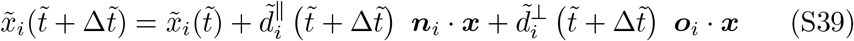

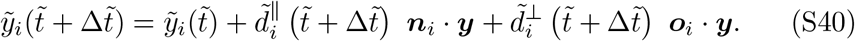
6. Update the orientation of cell *i*

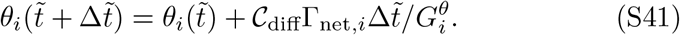

The ranges of values of the parameters used in our simulations are shown in Table S2.

We also conduct simulations at a smaller time step to verify that the simulation output is independent of the time step chosen, as shown in Fig. S3.

**Table S2:**
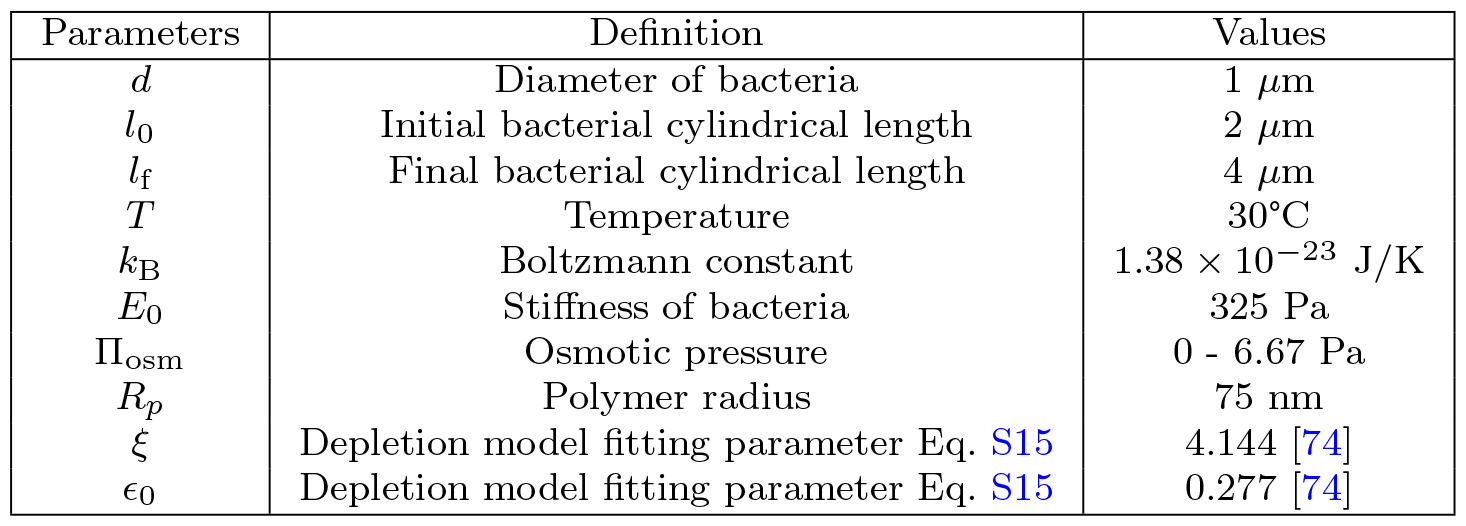
Parameters employed in agent-based simulations.

**Fig. S3:**
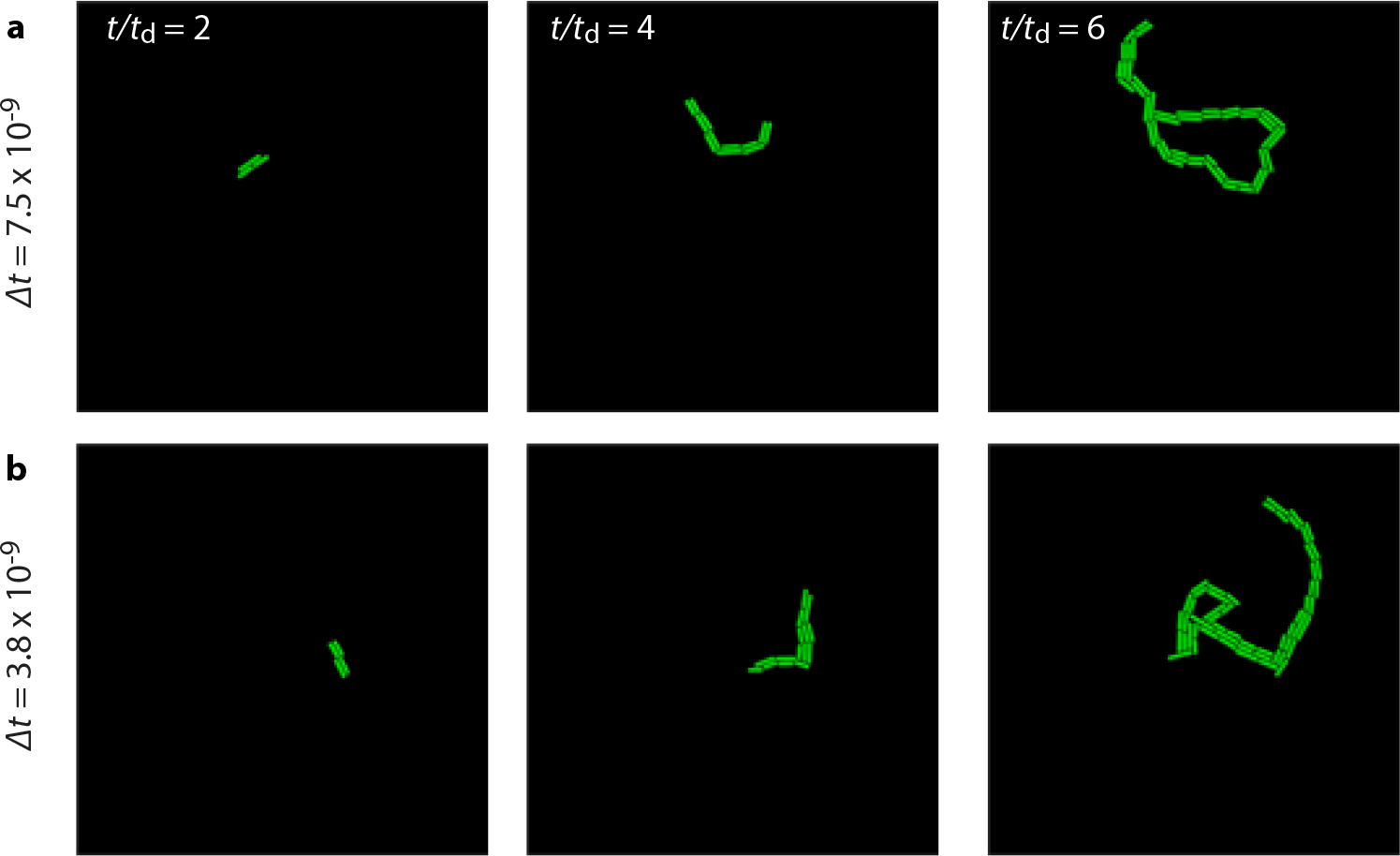
Smaller time step still leads to cable formation in our agent-based simulations. **a-b** Show time sequences of two representative simulations where we reduce the time step by a factor of two. Visual inspection suggests that, in our simulations, cable formation is independent of the time step implemented. Here *t*_rot_/*t*_d_ = 0.05 and *U*_ee_*/k*_B_*T* = 3.88.

### Identification of simulated cables using graph theory

For Fig. 5, to identify cables from the simulation results for Fig. 5, we use graph theory. In particular, since we have access to the centroids of the cells as well as their respective lengths and orientations, we plot graphs and compute the graph’s eccentricity via these steps:

1. Construct an undirected graph by taking each cell as a node in the graph. There exists an edge between two cells (or nodes) *i* and *j* only when cell *i* and *j* are in physical contact with each other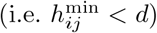. The length of said edge will be the shortest distance between the line segments of cells *i* and *j*.
2. We compute the graph’s eccentricity, *ℓ*, defined as the maximum distance amongst all the shortest paths between all nodes *i* and *j*. If said graph has multiple connected components we take the maximum eccentricity amongst all connected components.
3. For different colony morphologies we expect different graph eccentricities. For example, a random disconnected dispersion of cells yields *ℓ* = 0, while for *m* cells stacked side-by-side *ℓ* = *d*(*m* − 1). Given these different eccentricities for the growth morphologies we set an ad-hoc cut-off to distinguish between dispersed growth, stack growth, and cable growth (an example of these different graphs along with their eccentricities is shown in Fig. S4). A dispersed morphology was discerned for eccentricities *ℓ* < *d*, a stack morphology for eccentricities *d* < *ℓ* < 20*d*, and a cable morphology for eccentricities *ℓ >* 20*d*. The classification criteria is implemented when the simulation reaches the 21-27 cell stage because, at this growth stage, the three different morphologies can be visually discerned. Each simulation data point shown in Fig. 5a corresponds to an average over at least three individual simulations.

**Fig. S4:**
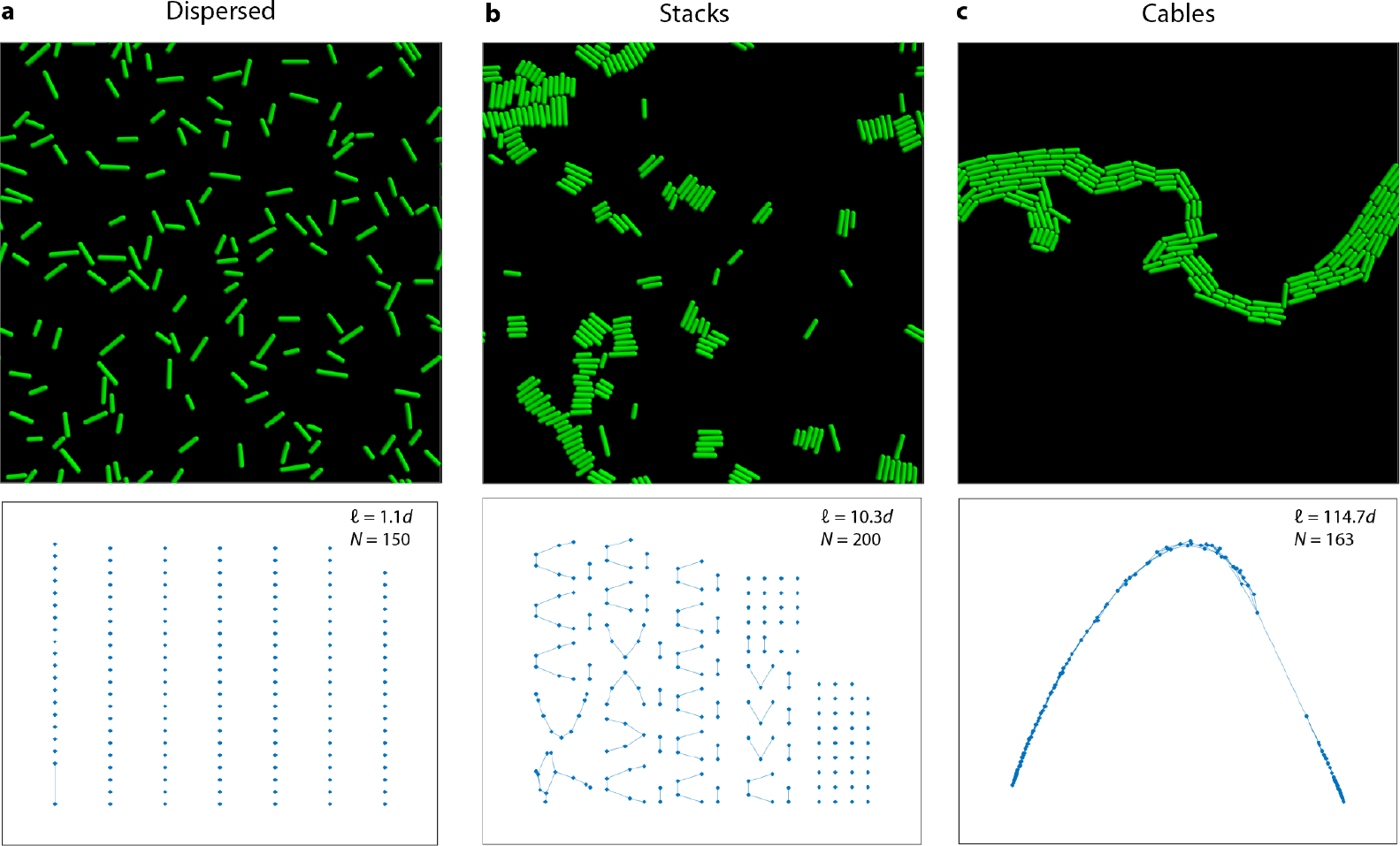
Graph eccentricity depends on growth morphology in simulations. **a-c**, Show representative snapshots of simulation outputs and their respective undirected graphs. The variables *N, ℓ*, and *d* correspond to the number of cells, the graph eccentricity, and the bacterial diameter respectively. **a**, Shows a snapshot of the Dispersed growth morphology characterized by cells that are mostly not in physical contact with one another, hence we observe a mostly disconnected graph with a small eccentricity *ℓ* = 1.1*d*. **b**, Shows a snapshot of the stack growth morphology characterized by cells that are aggregated in a side-by-side fashion, the formation of these stacks renders a more connected graph than shown in **a** hence it exhibits a higher graph eccentricity *ℓ* = 10.3*d*. **c**, Shows a snapshot of the cable growth morphology where now, all cells are connected in both a side-by-side and an end-to-end fashion, hence due the high level of connectivity of this morphology it exhibits the highest eccentricity *ℓ* = 114.7*d*.

### Calculation of the depletion potential between 4 cells

First, we compute the total depletion energy of *N* cells in general, only considering pairwise interactions:

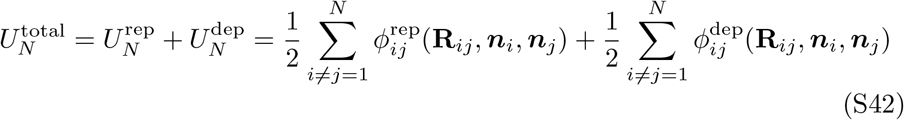

where *ϕ*_*ij*_ corresponds to the two types of pairwise interactions between cells: elastic repulsion and depletion attraction. For this calculation, for simplicity, we model the cells as infinitely hard objects, thus, the repulsive interaction potential has the form

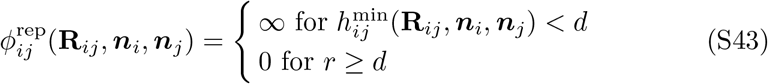

where 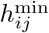 is the smallest distance between the centerlines of spherocylinders *i* and *j*. This quantity is itself a function of **R**_*ij*_, ***n***_*i*_, and ***n***_*j*_ corresponding to the vector joining the two center of masses of cells *i* and *j* and their orientation vectors respectively.

To calculate the total depletion energy of *N* cells we need to compute the total overlap volume of the system, *V*_*N*_. It has been shown that the total overlapping excluded volume between *N* bodies can be decomposed into zero-body, one-body, two-body, and higher-body terms, and that the higher-body terms can be neglected if the ratio between the polymer chain size and the cell diameter, *q*, does not exceed 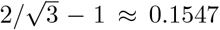 [74]. In our work, we employ a value of *q* = 0.15 because it is comparable to the polymer size used in experiments. Hence we can approximate the overlap volume between *N* cells as the sum of two-body contributions. To compute the total overlap volume, we sum over all *N* cells

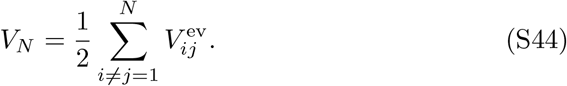

Equation S44 now enables us to calculate the depletion potential between 4 cells, 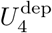, for the analysis of Fig. 4. In particular, to calculate the energy landscape shown in Fig. 4, we use a custom C++ script in which we consider 2-cell composites of 2 cells held side-by-side. One composite is held fixed at the origin with one cell at a coordinate (0, *d/*2) and another at (0, − *d/*2), where *d* is the cell’s diameter. The other composite is translated and rotated around the fixed composite, and we compute 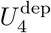 following the algorithm:

1. With one cell composite fixed at the origin we translate the center of mass of the movable composite to coordinate (*x*_CoM_, *y*_CoM_).
2. With fixed center-of-mass (*x*_CoM_, *y*_CoM_), we rotate the movable composite by 360° and as we rotate the composite we compute the total overlapping excluded volume, *V*_4_, associated with that angle of rotation. We only compute the total overlapping excluded volume if the smallest distance between any two cells in different cell composites, 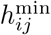, is bigger than *d* yet smaller than 2*R*_p_ + *d* where *R*_p_ is the polymer radius.
3. We obtain a list of overlapping excluded volumes at that particular (*x*_CoM_, *y*_CoM_) location corresponding to different orientations of the composite. Next we identify the largest overlapping excluded volume and take that as our estimated value of the overlapping excluded volume between 4 cells, *V*_4_. To simply characterize the depletion potential landscape, we assume that cells will always attain the configuration with the largest overlapping excluded volume at that specific (*x*_CoM_, *y*_CoM_) location. Hence with this information we can estimate 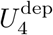 (*x*_CoM_, *y*_CoM_) as shown in Fig. 4b.

### Calculation of the duration 4 cells spend in the metastable interdigitated end-to-end configuration

To calculate *t*_esc_ in Fig. 5, we consider the two 2-cell composites discussed above with the following assumptions:

1. We model 4 cells as two 2-cell composites each consisting of 2 cells in a side-by-side configuration. One composite is fixed at the origin and its long and short axis span the space in which the second composite can rotate and translate.
2. For the purposes of estimating a diffusion coefficient for the movable 2-cell composite, *D*_2,rot_, we assume that the composite is an ellipsoid with one major axis equal to a cell’s long axis, *l*, and a minor axis equal to twice a cell diameter, 2*d*.
3. Visual inspection of the depletion energy landscape shown in Fig. 4b reveals a low-energy trajectory that the 2-cell composite can follow to escape the metastable interdigitated state (light blue curve with overlaid orange hollow circles in Fig. 4b). We term this trajectory the “minimal energy contour” and we assume that this is the only possible escape trajectory. This last assumption allows us to define a single reaction coordinate, *θ*, defined as the angle between the vector connecting the centroids of each 2-cell composite and the line spanned by the major-axis of the fixed 2-cell composite (Fig. 4b,c).

After plotting the configuration of 4 cells at different sites of the minimal energy contour we observe that the 2-cell composite rotates as it traverses the contour. Hence, for the following estimation, we will assume that rotation is the main motion that a 2-cell composite needs to perform in order to escape the depletion-induced energetic trap. Disregarding the translation of the 2-cell composite leads to an underestimate of the friction felt by the composite, thus our calculation yields an underestimate of the escape time. Lastly, to estimate the escape time of a 2-cell composite, at the 4-cell stage, we need to consider the height of the energetic barrier as well as its curvature, we take these two aspects into account by using Kramers’ escape formalism [76]

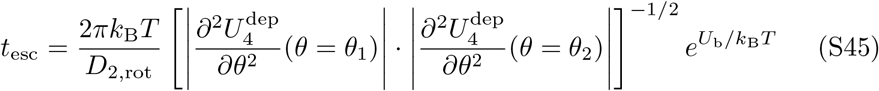

where 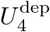 is the total depletion energy of 4 cells, *U*_b_ is the energetic barrier, *k*_B_ is the Boltzmann’s constant, *T* is temperature, *θ*_1_ corresponds to the angle where the local minimum is located, i.e. where 4 cells are in the interdigitated configuration, *θ*_2_ corresponds to the angle where the global minimum is located, i.e. where all 4 cells are in a side-by-side configuration, and the symbol |… | refers to an absolute value.

Eq. S45 allows us to relate the escape time to both the rotational diffusion of a 2-cell composite, *D*_2,rot_, and the end-to-end depletion energy of 2 cells. Indeed, we can measure the energy barrier *U*_b_ in units of *U*_ee_ and, furthermore, we can decompose the total depletion energy of 4 cells, 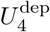, as the product between the osmotic pressure, Π_osm_, and the overlapping excluded volume, *V*_4_. With this decomposition we can isolate the geometrical shape of the potential profile and how this shape changes as the osmotic pressure varies and relate it back to *U*_ee_ because *U*_ee_ = Π_osm_*V*_ee_. We arrive at our final expression

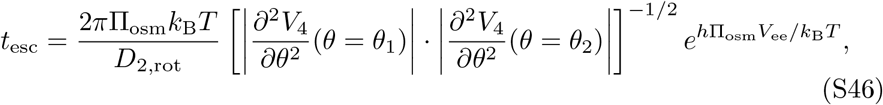

where *h* and *V*_ee_ are constants that depend on the polymer radius and bacterial diameter, while Π_osm_ changes, and thus modulates the value of the energetic barrier *U*_b_. This expression only holds for when *U*_b_ ≳ *k*_B_*T*. Notice that in this expression when Π_osm_ = 0, the escape time equal zero. Moreover, as Π_osm_ → ∞, the escape time also approaches infinity. Moreover, as *D*_2,rot_ → ∞, the escape time approaches zero as expected.

Having obtained this relation, we finally determine the transition boundary of Fig. 5 by setting *t*_esc_ = *t*_d_. In particular, rearranging Eq. S46 and multiplying both sides by *t*_2,rot_ = 1*/D*_2,rot_ as well as the ratio between the geometrical factors respective to 1-cell and 2-cell composite, 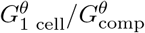, yields the final equation for the boundary shown in Fig. 5:

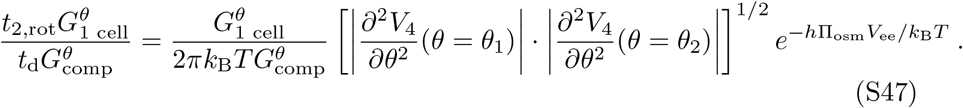

Importantly, this transition boundary does not strongly depend on the size ratio *q* = 2*R*_*p*_*/d*, as shown in Fig. S5.

**Fig. S5:**
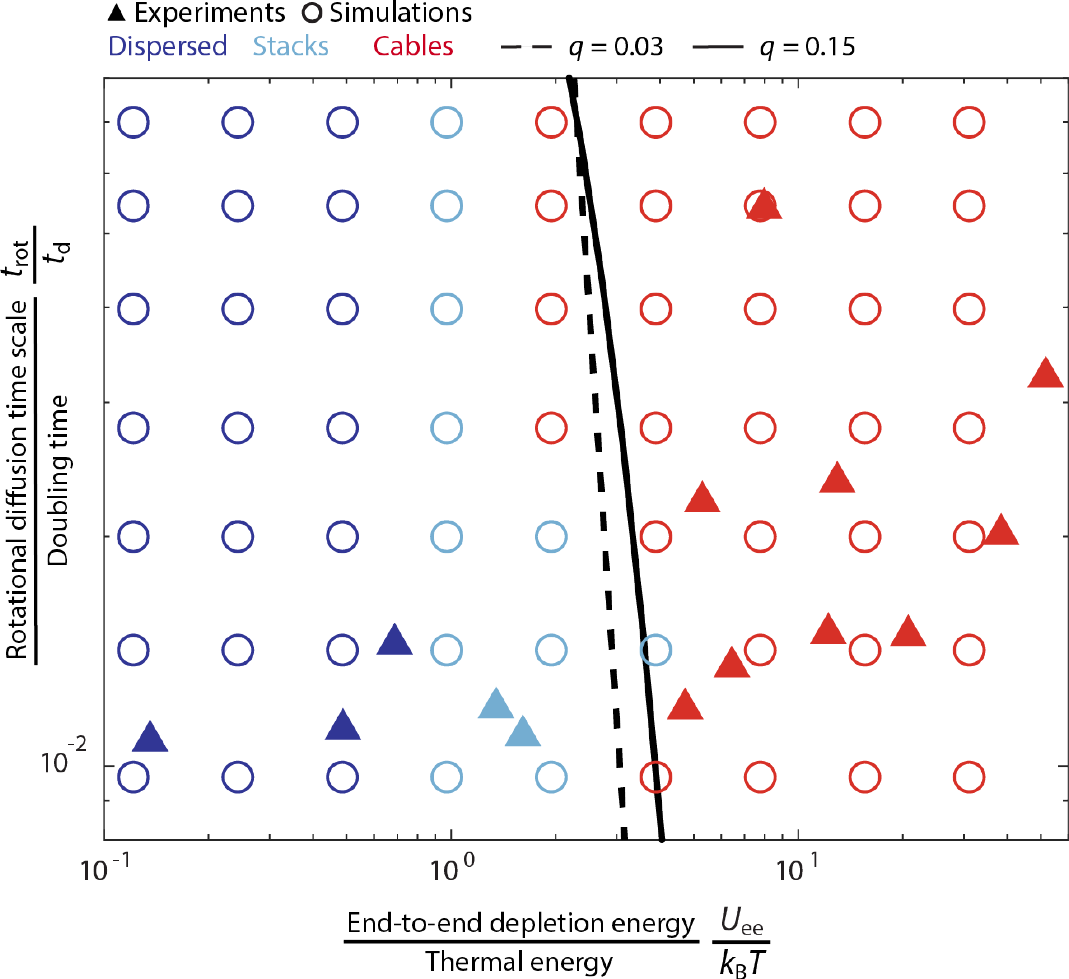
The model parameter *q* = 2*R*_p_*/d* does not significantly affect the transition boundary that delineates cable formation. Same figure as shown in the main text (Fig. 5a) but with two transition boundaries plotted for two distinct *q* values: 0.15 (solid) and 0.03 (dashed) corresponding to different polymer sizes.

### Physicochemical properties of mucus and how they influence cable formation

Where in Fig. 5 do biological polymers like mucus lie? We use literature reports of mucus mesh sizes Λ, viscosities *η*, and bacterial doubling times in mucus *t*_d_ (Table S3) to estimate the corresponding values of the two nondimensional parameters *U*_ee_*/k*_B_*T* and *t*_rot_*/t*_d_ in mucosal environments.

We estimate *U*_ee_*/k*_B_*T* as [71, 73]

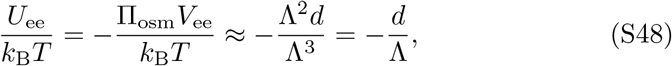

and estimate the diffusive time *t*_rot_*/t*_d_ using Eq. S29 described previously. These calculations yield the shaded grey region in Fig. 5a.

### Interplay between cellular motility and polymer-induced depletion interactions

As a first investigation of cable formation for *motile* cells, we repeat the experiment of Fig. 1, but with motile *E. coli* in LB media with 0.5 w/w% PEO 5 MDa. In this case, we do not observe cables (Movie S12).

Why does cellular motility appear to hinder cable formation? To gain insight into the underlying reason, we compare the hydrodynamic force *F*_prop_ = 3*πηdv* where *η* is the viscosity of the medium, *d* the bacterial diameter and *v* the swimming speed of *E. coli* cells [84], which enables them to overcome attractive forces, to the attractive depletion force induced by polymers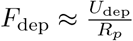.

For test polymer concentrations of 0.1 w/w% and 0.5 w/w% PEO 5 MDa, we use direct visualization of swimming cells to measure mean swimming velocities of 37 *µ*m/sec and 27 *µ*m/sec, respectively. Thus, *F*_dep_*/F*_prop_ = 0.2 to 0.5, indicating that the depletion attraction is insufficient to overcome propulsive forces. Our results thus suggest that cellular motility hinders cable formation. However, it could be that for other polymers, cells, and environmental conditions, *F*_dep_*/F*_prop_ *>* 1. Further investigating the interplay between cellular motility and polymer-induced cable formation will be an interesting direction for future research.

**Table S3:**
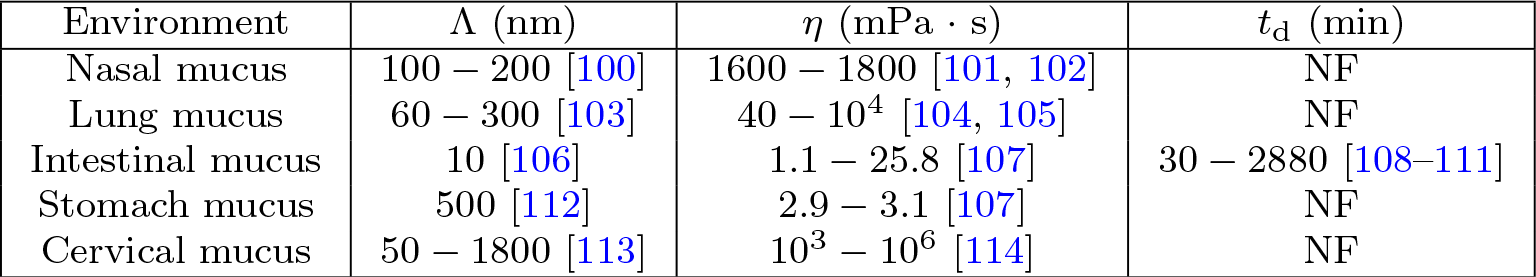
Ranges of literature values found for mucus mesh size (Λ), mucus viscosity (*η*), and bacteria doubling time (*t*_d_). Abbreviations: NF, not found.

## Additional supplementary figures referenced in the main text

**Fig. S6:**
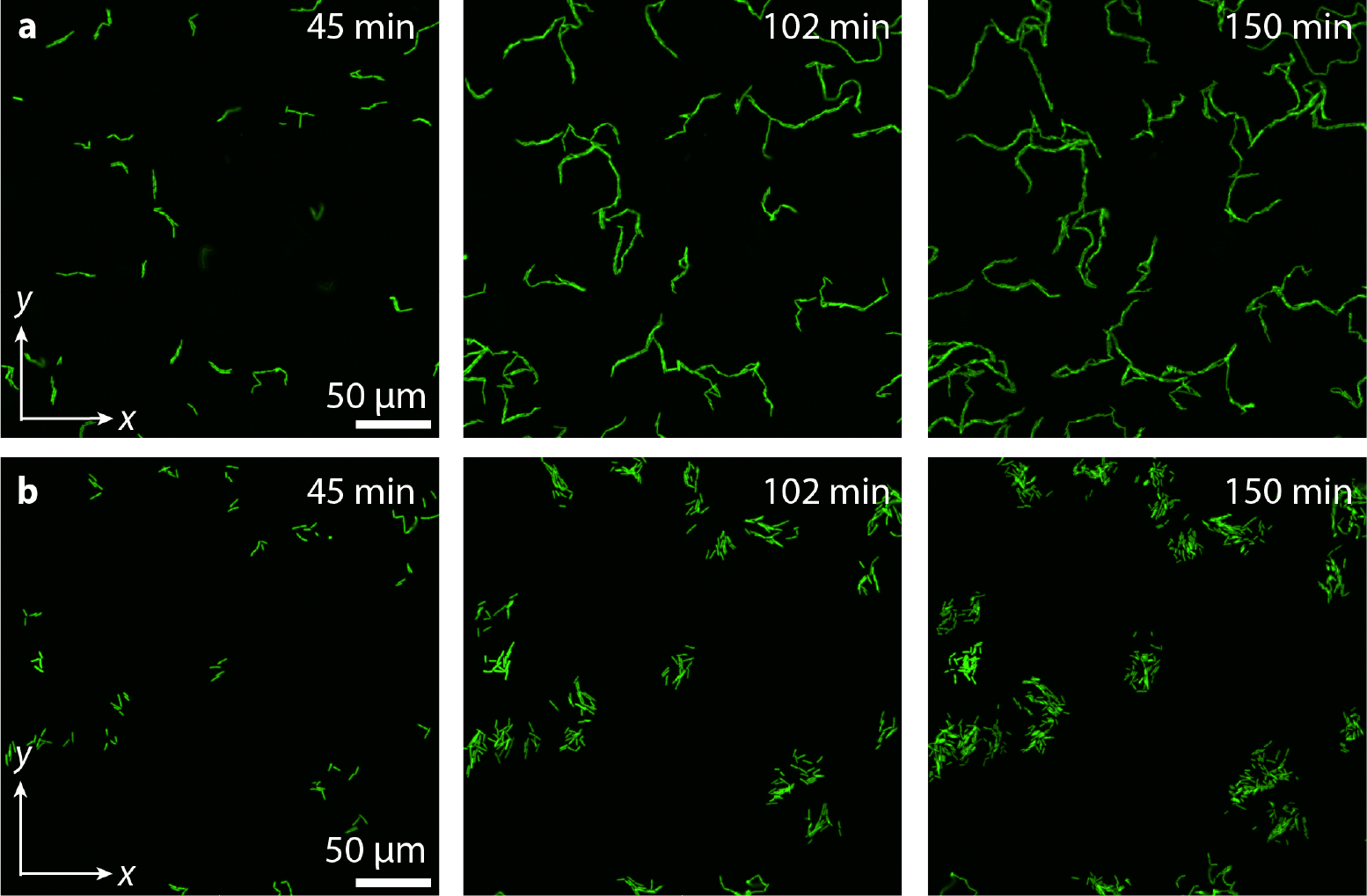
Non-motile *E. coli* cells grow into cable-like morphologies in the presence of Muc2 mucin polymers. **a**, Shows how *E. coli* cells at a mucin concentration of 0.5 wt% grow into cable-like morphologies. **b**, Same as in **a** but without any mucin polymer. The experiments were carried out in glass capillaries as described in the methods.

**Fig. S7:**
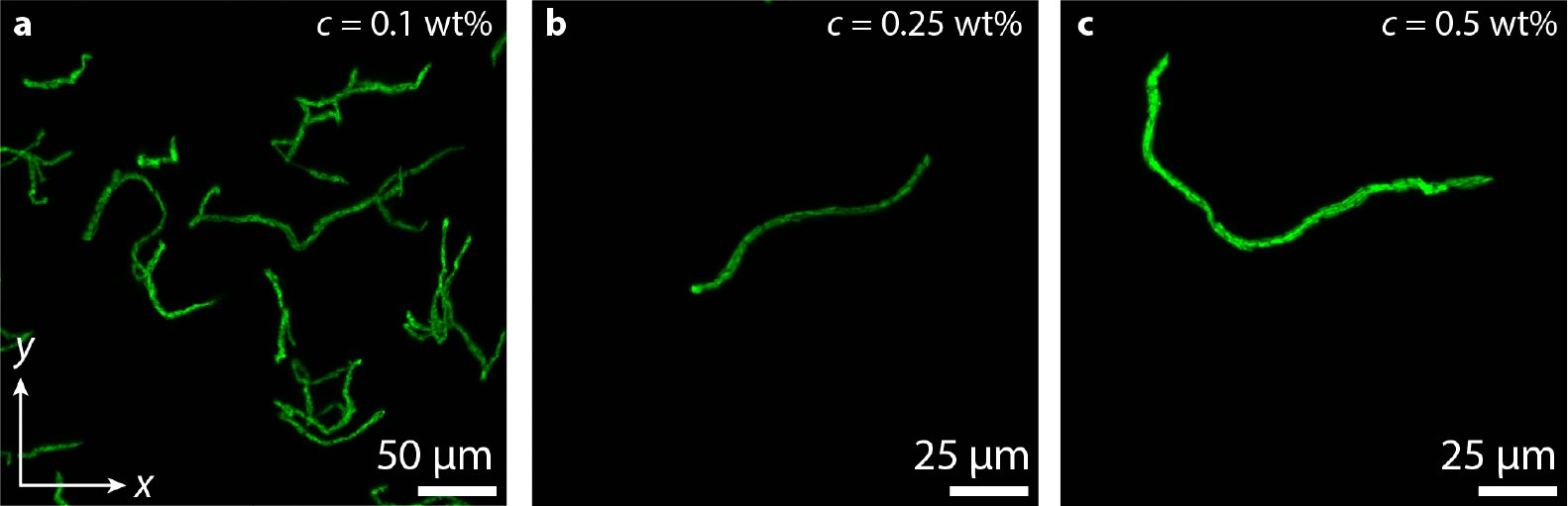
*E. coli* cable morphologies are observed in the bulk of polymer solutions away from solid boundaries. To grow and observe cable-like morphologies suspended in the bulk of a polymer solution we first grow immotile *E. coli* cells as described previously. Next, we mix cells in a polymer solution (PEO 5 MDa dissolved in LB at concentrations of **a**. 0.1 wt%, **b**. 0.25 wt%, and **c**. 0.5 wt%) and load them into a transparent-walled glassbottom Petri 35 mm in diameter. The imaging well of the dish is sealed with a PDMS slab to minimize evaporation. The dish is then fastened to a VWR tube rotator which is then kept inside a static 30°C incubator. The samples are left spinning inside the incubator for a minimum of three hours and thirty minutes. After this incubating period has passed we take the samples off the rotor and quickly image our samples in an A1R+ inverted laser-scanning confocal microscope with a 20x air objective. The microscope stage is maintained at 30 ± 1 °C. The three panels above show maximum intensity projection images of the cable morphologies thereby obtained. The images were taken at least 50 *µ*m above the glass bottom surface.

**Fig. S8:**
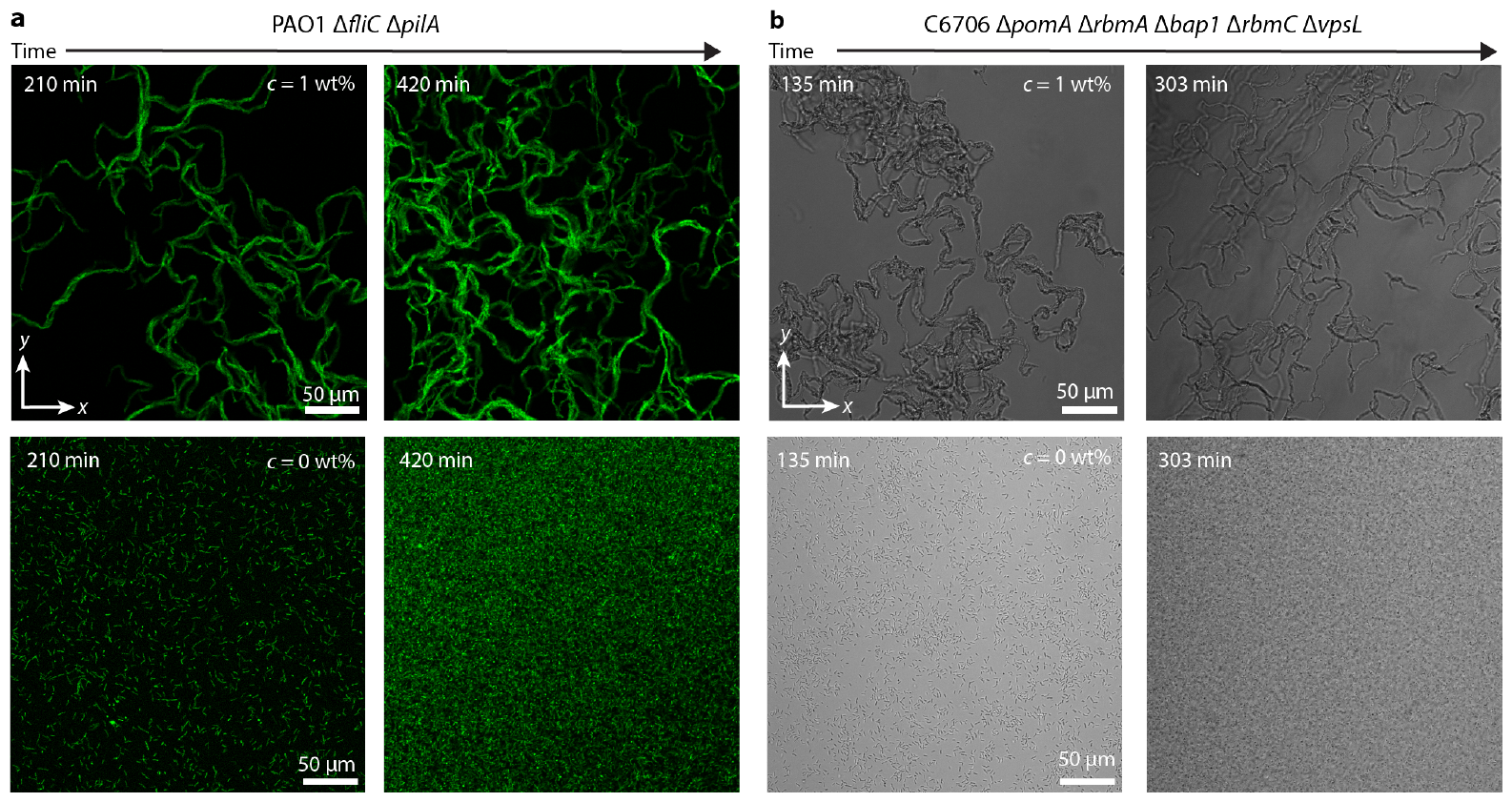
Non-motile strains of *P. aeruginosa* and *V. cholerae* form cable-like morphologies in the presence of polymers. **a**, Shows *P. aeruginosa* cells proliferating in both a polymer solution of 1 wt% PEO 1 MDa and without the presence of polymers. **b**, Same as in **a** but with *V. cholerae cells*. Each column corresponds to two different times where we have defined *t* = 0 min as the when the 96 well-plate is inserted into the static incubator to later image. The images were taken at different spatial locations of the same single-well at different time points.

**Fig. S9:**
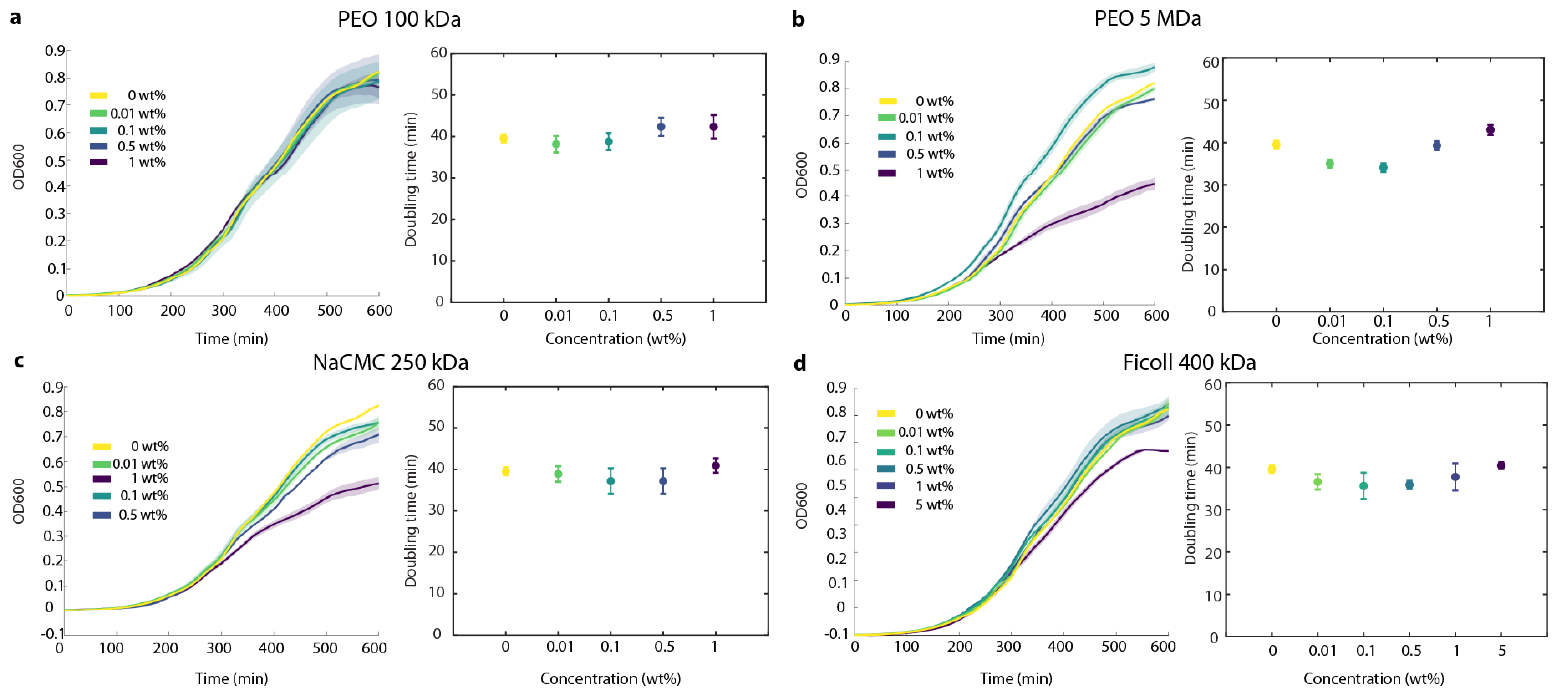
*E. coli* doubling times are similar across different polymer solutions. **a-d** Show both the growth curves and doubling times for *E. coli* cells in different polymer solutions. The growth curves were obtained in triplicate where the solid curve represents the mean among all samples while the shaded region is its corresponding standard deviation. The doubling times were measured by fitting the first 500 min of each sample’s growth curve to the Gompertz equation [115]: OD = *A* exp 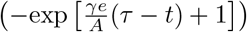 where *t* is time, *A* is the maximum value that the growth curve attains in the time window, *γ* is the maximal growth rate, *τ* is the lag time, and OD is the optical density (OD_600_). We convert the bacteria’s maximal growth rate, *γ*, to a doubling time via *t*_d_ = log(2)*/γ* which are reported in Table S1. We observe that, throughout all polymer concentrations tested, *E. coli* ‘s doubling time remains between 35-45 min and does not depend appreciably on polymer concentration. We obtained the cell’s growth curves under different polymer concentrations tested via optical density measurements in a Biotech Epoch 2 microplate reader. We inserted a 96 well plate into the plate reader where each well had 200 *µ*L of polymer solution and was initially seeded with 2 *µ*L of *E. coli* that was previously grown for 3 hours with 2 wt% LB. The optical density readings were recorded every 15 min at 30°C with the 96 well plate continually shaking at a frequency of 597 cpm and amplitude of 3 mm.

**Fig. S10:**
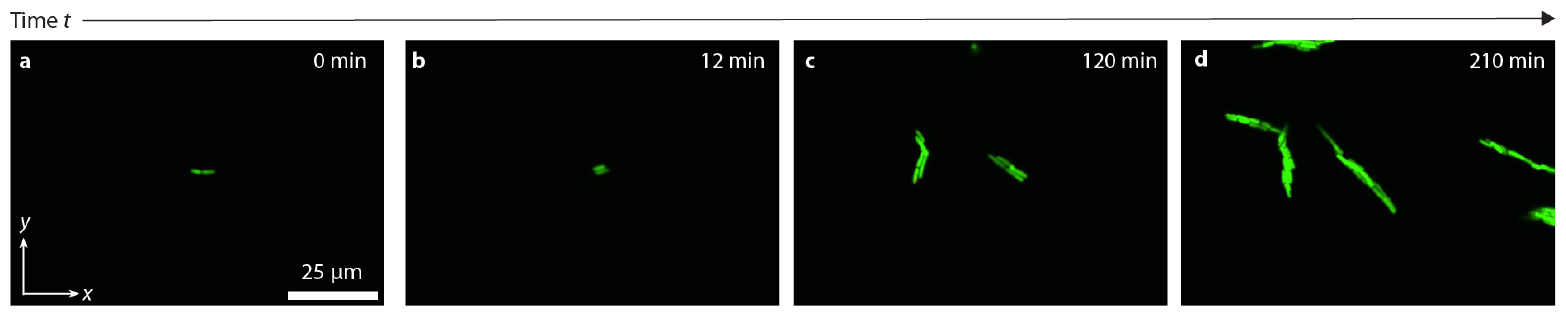
Early cable dynamics of *E. coli* cable morphologies. The four panels above show a time sequence of images of cable morphologies in polymer solution of 0.2 w/w% PEO 5 MDa from Movie S8. **a**, Pair of cells oriented end-to-end. **b**, Pair of cells quickly lock in side-by-side. **c**, Cells continue to proliferate end-to-end **d**, Cells create a cable morphology.

**Fig. S11:**
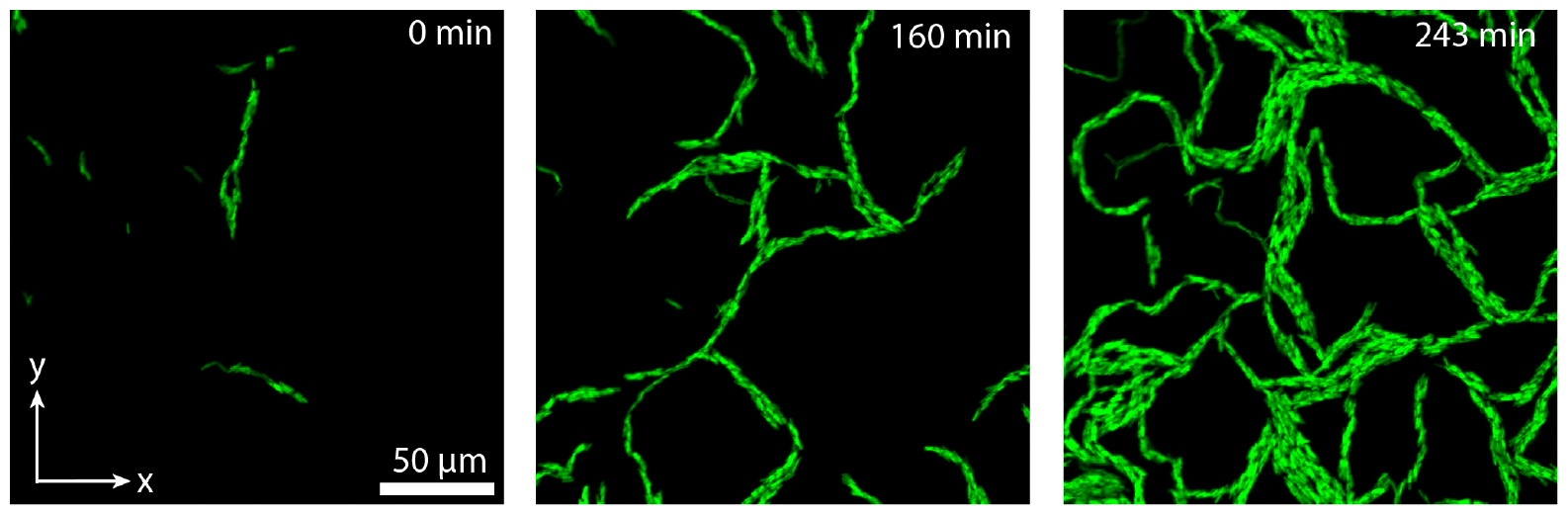
Cable-like morphologies do not break due to pump-induce shear stresses. Time sequence images of *E. coli* cells proliferating in the presence of 0.1 wt% PEO 100 kDa while, simultaneously, polymer solution at the same polymer concentration is pumped at a flow rate of 5 *µ*L*/*min (equal to that used in Fig. 2b). We define *t* = 0 as the time when pumping starts.

## Notes

https://zenodo.org/doi/10.5281/zenodo.10966670

### Competing Interest Statement

The authors have declared no competing interest.

